# Bacterial Efflux Pump Modulators Prevent Bacterial Growth in Macrophages and Under Broth Conditions that Mimic the Host Environment

**DOI:** 10.1101/2023.09.20.558466

**Authors:** Samual C. Allgood, Chih-Chia Su, Amy L. Crooks, Christian T. Meyer, Bojun Zhou, Meredith D. Betterton, Michael R. Barbachyn, Edward W. Yu, Corrella S. Detweiler

## Abstract

New approaches for combatting microbial infections are needed. One strategy for disrupting pathogenesis involves developing compounds that interfere with bacterial virulence. A critical molecular determinant of virulence for Gram-negative bacteria are efflux pumps of the resistance-nodulation-division (RND) family, which includes AcrAB-TolC. We previously identified small molecules that bind AcrB, inhibit AcrAB-TolC, and do not appear to damage membranes. These efflux pump modulators (EPMs) were discovered in an in-cell screening platform called SAFIRE (Screen for Anti-infectives using Fluorescence microscopy of IntracellulaR Enterobacteriaceae). SAFIRE identifies compounds that disrupt the growth of a Gram-negative human pathogen, *Salmonella enterica* serotype Typhimurium (*S.* Typhimurium) in macrophages. We used medicinal chemistry to iteratively design ∼200 EPM35 analogs and test them for activity in SAFIRE, generating compounds with nanomolar potency. Analogs were demonstrated to bind AcrB in a substrate binding pocket by cryo-electron microscopy (cryo-EM). Despite having amphipathic structures, the EPM analogs do not disrupt membrane voltage, as monitored by FtsZ localization to the cell septum. The EPM analogs had little effect on bacterial growth in standard Mueller Hinton Broth. However, under broth conditions that mimic the micro-environment of the macrophage phagosome, *acrAB* is required for growth, the EPM analogs are bacteriostatic, and increase the potency of antibiotics. These data suggest that under macrophage-like conditions the EPM analogs prevent the export of a toxic bacterial metabolite(s) through AcrAB-TolC. Thus, compounds that bind AcrB could disrupt infection by specifically interfering with the export of bacterial toxic metabolites, host defense factors, and/or antibiotics.

**IMPORTANCE:** Bacterial efflux pumps are critical for resistance to antibiotics and for virulence. We previously identified small molecules that inhibit efflux pumps (efflux pump modulators, EPMs) and prevent pathogen replication in host cells. Here we used medicinal chemistry to increase the activity of the EPMs against pathogens in cells into the nanomolar range. We show by cryo-electron microscopy that these EPMs bind an efflux pump subunit. In broth culture, the EPMs increase the potency (activity), but not the efficacy (maximum effect), of antibiotics. We also found that bacterial exposure to the EPMs appear to enable the accumulation of a toxic metabolite that would otherwise be exported by efflux pumps. Thus, inhibitors of bacterial efflux pumps could interfere with infection not only by potentiating antibiotics, but also by allowing toxic waste products to accumulate within bacteria, providing an explanation for why efflux pumps are needed for virulence in the absence of antibiotics.

## INTRODUCTION

Infections caused by Gram-negative bacteria are particularly challenging to treat because their cell envelope incorporates complementary defenses that protect them from chemicals (1–3). The outer membrane has an external leaflet that is relatively impermeable to chemicals because it consists of tightly packed, negatively charged lipopolysaccharide (LPS) molecules bound to magnesium cations, which together repel hydrophobic molecules (4). The outer membrane surrounds a porous cell wall and the inner membrane. Compounds that manage to breach the outer membrane are typically captured in the periplasm or inner membrane by efflux pumps and are exported across both membranes. Thus, antibiotics that slowly permeate the outer membrane are prevented from accumulating to critical levels within the bacterium (2, 5, 6).

Select efflux pumps are demonstrated virulence determinants, meaning they are required for bacteria to cause infection even in the absence of treatment with antibiotics (7–10). This observation could reflect that efflux pumps are needed to expel metabolic waste and/or host defense molecules during infection. Nevertheless, these same efflux pumps are not essential for bacterial growth in standard broth culture (6). In patients treated with commonly used antibiotics, bacterial efflux pumps typically contribute to bacterial multi-drug resistance (MDR) by capturing and exporting the antibiotics, increasing the probability of treatment failure and the emergence of resistant clones (11). For these reasons, there is interest in developing inhibitors of efflux pumps that could re-sensitize MDR bacteria to antibiotics and/or prevent pathogen export of toxic metabolites and host defense molecules (6, 12).

In *E. coli* and the closely related *Salmonella enterica* serotype Typhimurium (*S.* Typhimurium) the major efflux pump required for virulence is AcrAB-TolC, a member of the classic resistance-nodulation-division (RND) family. AcrAB-TolC is a multi-subunit machine that spans the inner membrane, periplasm, and outer membrane (13). In the inner membrane, AcrB assembles into a trimer and traps substrates from the lipid bilayer or the periplasm. The AcrB trimer has a large substrate binding pocket that can manage substrates with diverse structures. AcrB transfers substrates to the periplasmic subunit of the pump, AcrA, which in turn passes substrates to the TolC channel in the outer membrane, enabling extrusion out of the cell (14–17).

We previously described three small molecules that inhibit AcrAB-TolC and were called efflux pump modulators (EPMs): EPM30, EPM35 and EPM43. These EPMs were discovered in a screen for compounds that prevent *S.* Typhimurium from colonizing macrophages in a screening platform called SAFIRE (Screen for Anti-infectives using Fluorescence microscopy of IntracellulaR Enterobacteriaceae) (18, 19). In the test tube, the EPMs bind AcrB, and in bacteria, they interfere with the export of substrates by AcrAB-TolC. In macrophages, the EPMs synergize with antibiotics that are exported by AcrAB-TolC, including ciprofloxacin, erythromycin and chloramphenicol suggesting they target AcrAB-TolC during infection (13, 18, 20, 21). The EPMs may also target other efflux pumps, as they synergize with cationic antimicrobial peptides not known to be AcrAB-TolC substrates in *E. coli,* such as LL-37 and polymyxin B (18, 22, 23). Moreover, the EPMs inhibit the survival of both wild-type and *acrAB-*deficient *S.* Typhimurium in macrophages and in HeLa cells. The EPMs did not have apparent deleterious effects on bacterial or mammalian membranes, a key characteristic because dissipation of proton motive force deactivates efflux pumps non-specifically by depriving them of energy. However, these three EPMs disrupted mammalian cell morphology, indicating toxicity (18). We therefore embarked on an iterative approach that combined medicinal chemistry with activity screening in SAFIRE to improve EPM potency and decrease toxicity.

## RESULTS

### Design and Screening of EPM Analogs

All three of the EPMs previously identified were used to inform early structure-activity relationship (SAR) efforts because their activity profiles suggested distinct efflux modulation profiles (18). However, the three EPMs share related chemical motifs. For example, EPM30 contains a guanidine moiety, which is chemically similar to the aminopyrimidine moiety in EPM35 and the diamino-benzopyrimidine moiety in EPM43. We designed and synthesized analogs in batches of 20-50, up to approximately 200 chemical compounds. Our initial synthetic efforts primarily focused on varying the left-hand side, the right-hand side, and the diamine moieties of EPM35, with modifications to the 2-hydroxypropanyl central linker limited to simple O-alkylation and preparation of enantiomeric pairs to explore the effect of absolute configuration on activity (**Figure 1 A**). On the left-hand side, we examined analogs containing substitutions of the phenyl ring with, for instance, methoxy, alkyl, cyano, or trifluoromethyl groups, halogens, and several combinations thereof. On the right-hand side, we experimented with removing the 6-trifluoropyrimidin-2-yl group, consistent with another goal of the medicinal chemistry campaign, which was to identify the minimum pharmacophore for this series of compounds. We also altered the diamine subunits with 1) the introduction of N-alkyl, N,N-dialkyl and N-acetyl groups in place of the simple primary amine, as well as 2) investigating a variety of isosteric cyclic amine ring systems, for example pyrrolidine, to replace the starting piperidine ring of EPM35. Each of the 200 analogs was vetted by LC-MS for purity >95% and by ^1^H NMR for the intended structure.

**Figure 1.**
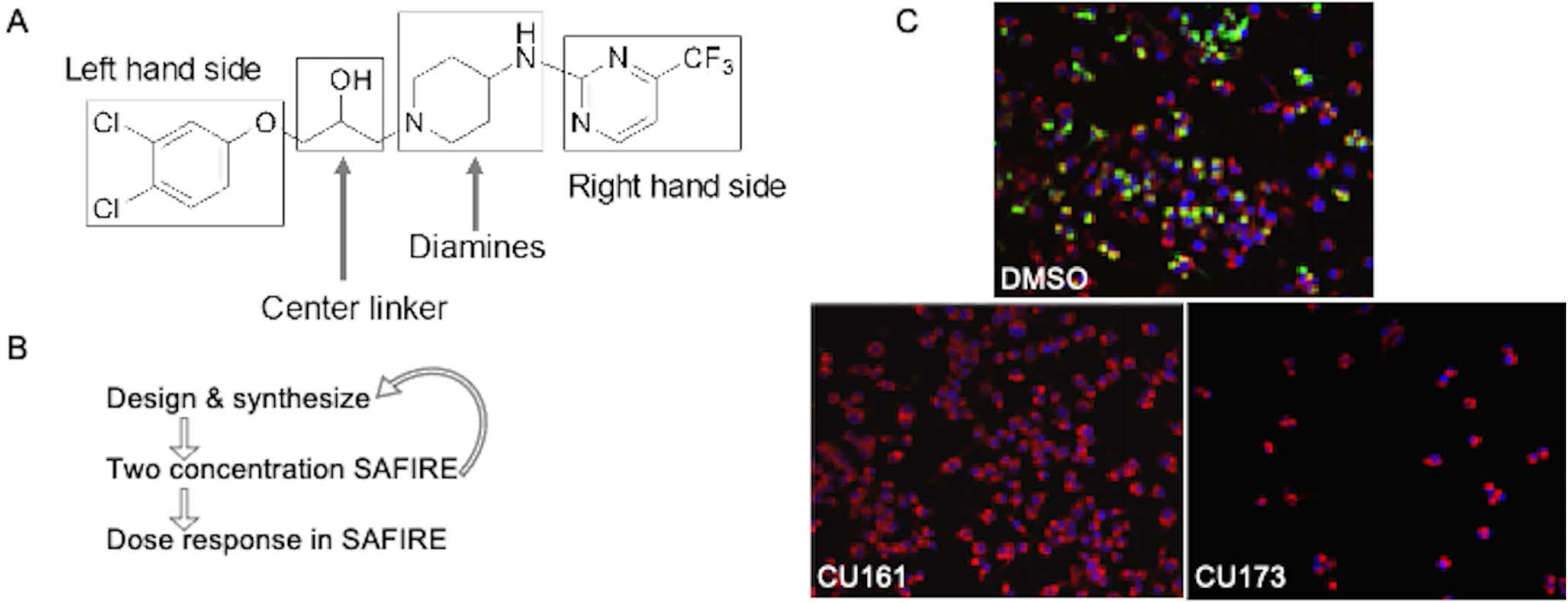
Approach for the improvement of EPM activity in the SAFIRE assay and reduction of toxicity. **A)** Structural components of EPM35 modified by medicinal chemistry. **B)** Iterative screening pathway. The two concentrations tested in SAFIRE were 5 and 25 μM. Dose responses were performed with at least 8 analog concentrations. **C)** Examples of infected macrophages treated with control (DMSO), a non-toxic (CU161) and a toxic (CU173) analog, as evidenced by the small number of macrophages apparent 16 hours after treatment. *sifB::gfp* (green), MitoTracker (red), DAPI (blue).

Analogs were iteratively tested for potency and toxicity at two concentrations, 5 and 25 µM, in the SAFIRE macrophage assay (**Figure 1B**). In SAFIRE, we used MATLAB software to count the number of macrophages that remain adherent and retain the red fluorescence of MitoTracker, an indicator of voltage across the mitochondrial inner membrane. MATLAB then calculates the signal from bacteria (GFP) that overlaps with macrophages, as defined by MitoTracker, to estimate bacterial load. This process revealed that substitutions on the amine nitrogen compromised antibacterial activity in SAFIRE (**Table S1**). If a compound reduced the percentage of adherent macrophages to 60% or less than DMSO-controls or resulted in poor macrophage morphology after 16 hours of exposure, then we eliminated the compound as host-toxic (**Figure 1C, Table S1**). For example, we observed that the 6-trifluoropyrimidin-2-yl group on the right-hand side of EPM35 is apparently toxic to macrophages. Non-toxic analogs that reduced bacterial load by at least 60% at 5 and 25 µM in SAFIRE, as compared with DMSO, progressed and were evaluated in SAFIRE across eight or more concentrations, ranging from 0.001 to 50 µM (n=2). Calculated IC_50_ values suggest that in-cell antibacterial activity increased more than 100-fold for some of the analogs compared to the parent compound EPM35 (**Figure 2**). Each of the top 13 analogs had at least an 8-fold increase in potency and were not toxic for macrophages at 50 µM in the SAFIRE assay (**Figure 1C**).

**Figure 2.**
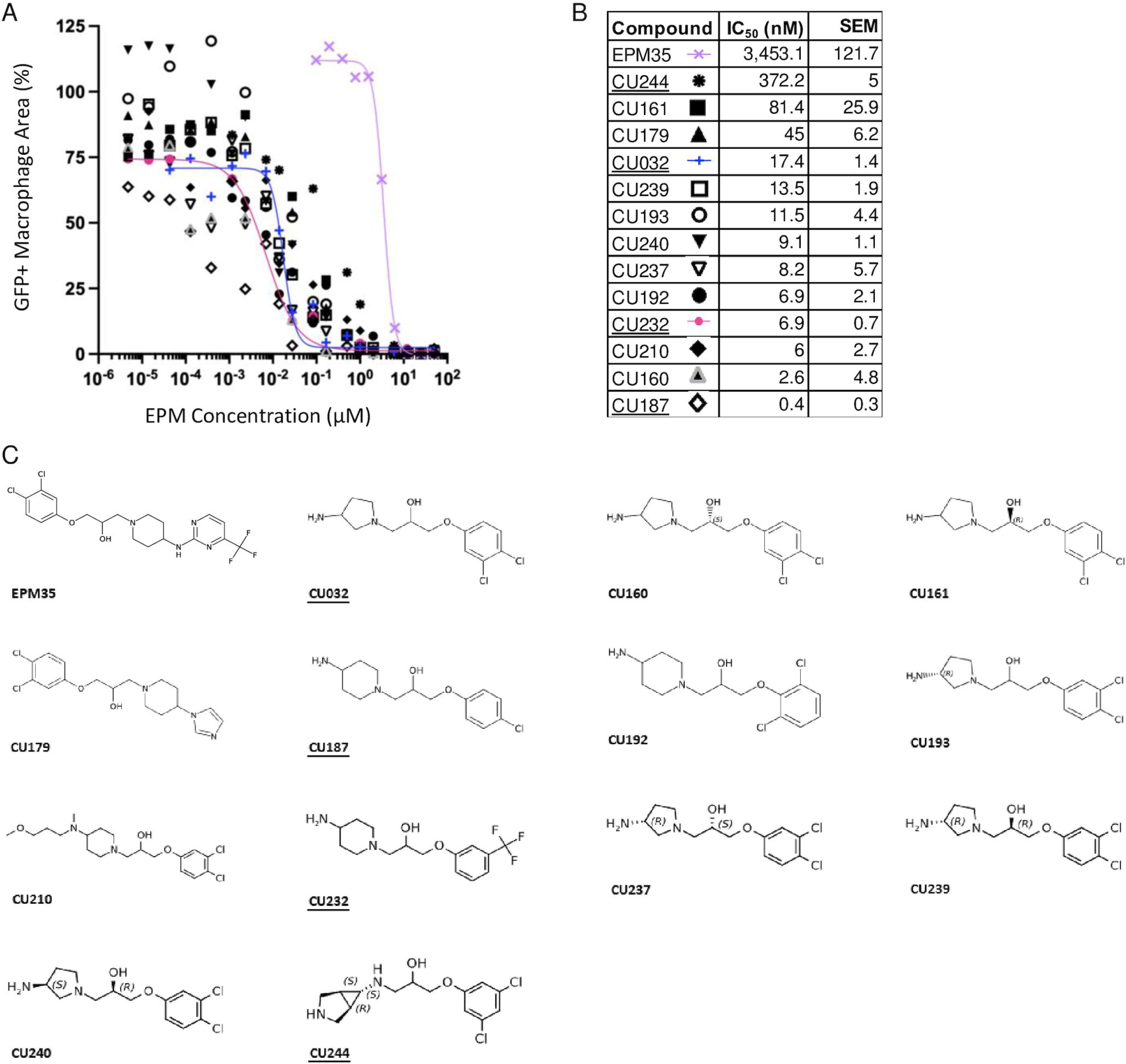
SAFIRE IC_50_ curves, values, and structures of EPM35 (parent compound) and the top 13 EPM analogs. **A, B)** IC_50_ curves and values. Data are derived from ≥ 10 concentrations per analog and are normalized to DMSO-treated control samples (100%). **C)** Structures. EPM analogs examined in subsequent figures are <u>underlined</u>.

### The EPM Analogs Interact with AcrB

We selected three compounds for further study, CU032, CU232, and CU244 because they incorporated different aspects of the parent structure, specifically, the negative atoms on the benzene ring and the amine on opposite side of the molecule. To determine whether the EPMs could bind AcrB, we used purified *E. coli* AcrB, which is 95% identical to *S. enterica* Typhimurium AcrB. We previously used isothermal titration calorimetry (ITC)(18, 24) to show that AcrB interacts with the EPM35 parent compound with a K_D_ in the sub-micromolar range (18). Here we show with ITC that, like EPM35, the three analogs are capable of specifically interacting with AcrB to form complexes (**Figure S1, Table 1**). Curiously, the analogs have K_D_s that are 25-50 -fold larger than that of EPM35. In addition, the binding of CU032 and CU232 to AcrB releases significantly less enthalpy (H) and increases entropy (S), indicating that these two EPM analogs rely on the entropic effect for binding, whereas binding of the parent EPM35 ligand to AcrB is more dependent on the enthalpic effect. Nevertheless, our ITC data revealed equilibrium dissociation constants (K_D_s) for AcrB and the EPM analogs that are in good agreement with those of AcrB substrates: K_D_s of AcrB with the EPM analogs range from 7.4 to 15 μM, whereas K_D_s of AcrB with rhodamine 6G, proflavine, and ciprofloxacin range from 5.5 to 74.1 µM (25). Thus, the EPM analogs interact specifically with AcrB in a manner consistent with established EPM substrates.

**Table 1.** Binding of EPMs to AcrB, as Calculated From ITC Results.

| Compound | $K_D$ (μM) | $\Delta H$ cal/mol | $\Delta S$ cal/(mol deg) |
| --- | --- | --- | --- |
| **EPM35<sup>1</sup>** | $0.29 \pm 0.03$ | $-7663 \pm 84$ | 4.2 |
| **CU032** | $14.7 \pm 9.56$ | $-1947 \pm 490$ | 15.60 |
| **CU232** | $7.41 \pm 1.00$ | $-2124 \pm 93$ | 16.30 |
| **CU244** | $15.0 \pm 3.81$ | $-8654 \pm 1384$ | -6.91 |

<sup>1</sup> EPM35 was previously reported (18) and is included here for comparison.

### EPM Analogs Bind to AcrB in the Substrate Hydrophobic Pocket

To elucidate at high resolution how the EPM analogs interact with AcrB, we used cryo-EM. We overproduced and purified the *E. coli* AcrB protein containing a 6xHis tag at the C-terminus and reconstituted the protein into lipidic nanodiscs (17). We incubated the AcrB-nanodisc samples with the parent compound or with one of three structurally dissimilar EPMs analogs (CU032, CU232, CU244), to form AcrB-EPM complexes. We then solved single-particle cryo-electron microscopy (cryo-EM) structures of these complexes to resolutions between 2.63 and 2.82 Å (**Table 2**, **Figure 3**).

**Figure 3.**
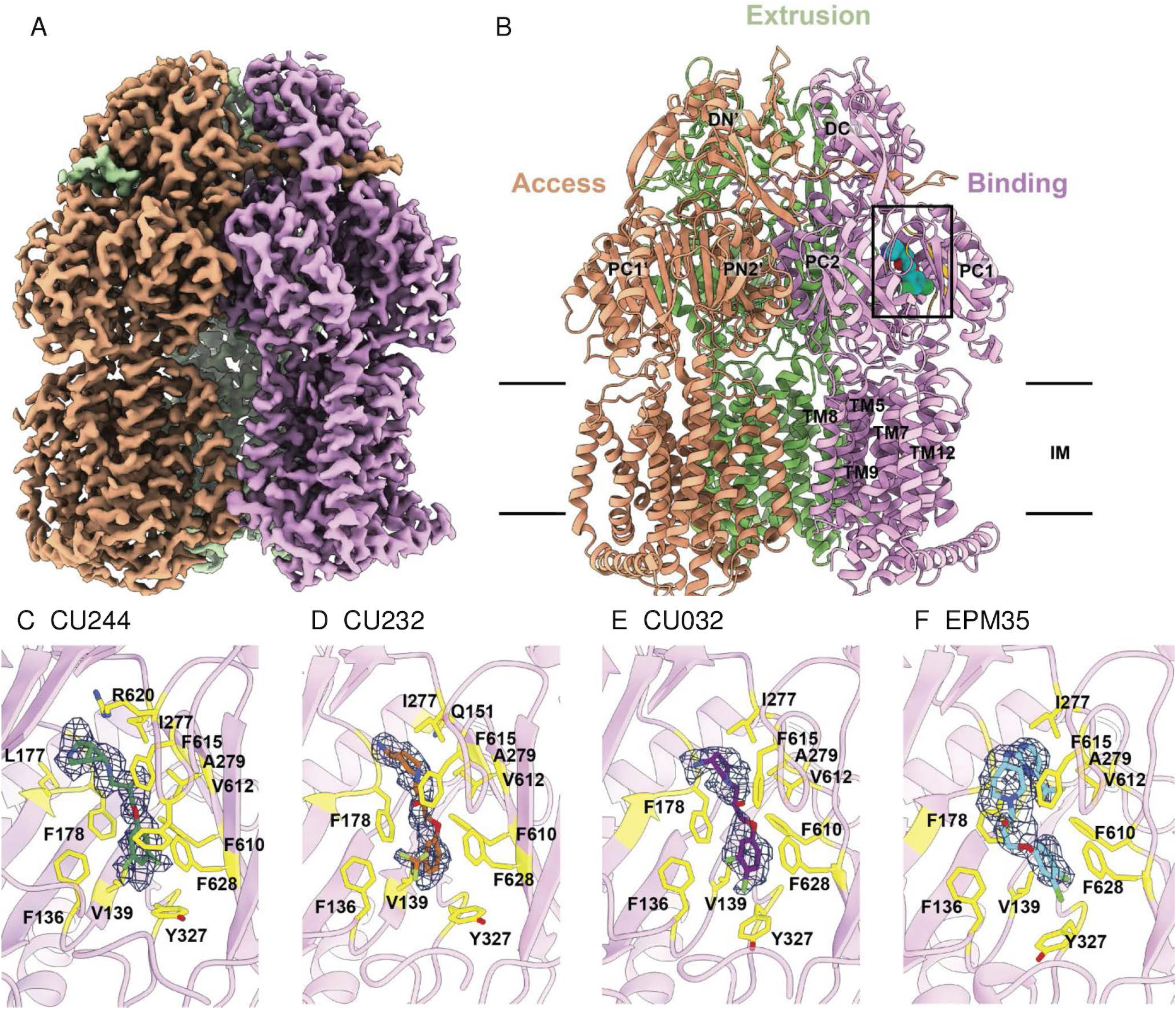
Single-particle cryo-EM structure of the AcrB-analog complexes. **A)** Cryo-EM density map of trimeric AcrB-CU244. **B)** Ribbon diagram of the structure of the side view of the AcrB-CU244 trimer. The bound CU244 molecule within the “binding” protomer of AcrB-CU244 is represented by cyan spheres. In (A-B), the “access”, “extrusion” and “binding” protomers are colored orange, green and pink, respectively. Each protomer of AcrB contains 12 transmembrane helices (TM1-TM12) and six periplasmic subdomains (PN1, PN2, PC1, PC2, DN and DC). **C)** The CU244 binding site. The bound CU244 molecule is colored green. **D)** The CU232 binding site. The bound CU232 molecule is colored orange. **E)** The CU032 binding site. The bound CU032 molecule is colored purple. **F)** The CU035 binding site. The bound EPM35 molecule is colored cyan. In (C-F), Residues participating in compound binding are in yellow sticks. The cryo-EM densities of bound compounds are colored dark blue.

**Table 2.** Cryo-EM data collection, processing, and refinement statistics.

| **Data Set** | **CU32** | **EPM35** | **CU232** | **CU244** |
| --- | --- | --- | --- | --- |
| **Data collection and processing** |  |  |  |  |
| Magnification |  | 81,000 |  |  |
| Voltage (kV) |  | 300 |  |  |
| Electron Microscope |  | Krios-GIF-K3 |  |  |
| Defocus range (μm) |  | -1.0 to -2.5 |  |  |
| Total exposure time (s) |  | 3.8 |  |  |
| Energy filter width (eV) |  | 20 |  |  |
| Pixel size (Å) |  | 1.07 |  |  |
| Total dose (e<sup>-</sup>/Å<sup>2</sup>) |  | 36 |  |  |
| Number of frames |  | 38 |  |  |
| Dose rate (e<sup>-</sup>/phys. pixel/s) |  | 16 |  |  |
| No. of initial micrographs | 2,683 | 2,229 | 1,267 | 1,002 |
| No. of initial particles | 1,680,995 | 3,244,867 | 1,270,215 | 998,124 |
| No. of final particles | 237,148 | 155,685 | 161,598 | 162,5048 |
| Symmetry | C1 | C1 | C1 | C1 |
| Resolution (Å) | 2.62 | 2.71 | 2.71 | 2.44 |
| FSC threshold | 0.143 | 0.143 | 0.143 | 0.143 |
| Resolution range | 2.39-6.79 | 2.50-9.80 | 2.40-7.15 | 2.39-9.84 |
| **Refinement** |  |  |  |  |
| Model resolution cut-off (Å) | 2.62 | 2.71 | 2.71 | 2.44 |
| **Model composition** |  |  |  |  |
| No. of Protein residues | 3,099 | 3,099 | 3,099 | 3,099 |
| **RMSD<sup>a</sup>** |  |  |  |  |
| Bond lengths (Å) | 0.003 | 0.003 | 0.003 | 0.004 |
| Bond angles (°) | 0.554 | 0.746 | 0.554 | 0.658 |
| **Validation** |  |  |  |  |
| MolProbity score | 1.13 | 1.42 | 1.13 | 1.22 |
| Clash score | 3.42 | 4.20 | 3.42 | 2.05 |
| **Ramachandran plot (%)** |  |  |  |  |
| Favored (%) | 98.16 | 96.57 | 98.16 | 99.03 |
| Allowed (%) | 1.84 | 3.43 | 1.84 | 0.97 |
| Disallowed (%) | 0 | 0 | 0 | 0 |
| CC mask | 0.82 | 0.81 | 0.82 | 0.83 |
| CC box | 0.66 | 0.65 | 0.64 | 0.65 |
| CC vol | 0.78 | 0.78 | 0.78 | 0.81 |

<sup>a</sup>, root mean square deviation

### Structure of AcrB-CU244

The single-particle images of the AcrB-CU244 complex led to a high quality cryo-EM map, allowing us to solve the structure at a resolution of 2.44 Å (**Table 2, Figure S2**). CU244 is housed in a large cavity created by the distal drug-binding site of the “binding” protomer of AcrB, where the periplasmic cleft is completely open. There are at least 23 amino acids involved in forming the distal drug-binding site of AcrB (26, 27). In addition, this distal site includes a hydrophobic trap that contributes strongly to drug binding (27). This trap includes residues F178, I277, V612 and F615. Within 4 Å of bound CU244, there are 12 residues, including F136, V139, L177, F178, I277, A279, Y327, F610, V612, F615, R620 and F628, involved in anchoring this EPM (**Figure 3C**). Most of these residues are hydrophobic in nature, suggesting that the binding is mainly governed by hydrophobic and aromatic interactions. Surprisingly, no hydrogen bonds are found to facilitate CU244 binding, highlighting the importance of these hydrophobic and aromatic residues for substrate recognition.

### Structure of AcrB-CU232

We incubated the AcrB-nanodisc sample with CU232 to form the AcrB-CU232 complex and solved this complex structure using single-particle cryo-EM. The reconstituted sample led to a cryo-EM map at a nominal resolution of 2.71 Å (**Table 2**, **Figure S3**). Superimposition of the AcrB-CU244 and AcrB-CU232 trimers gives rise to a calculated r.m.s.d. of 0.5 Å, indicating that the conformations of these two trimers are nearly identical to each other. Again, CU232 is only observed to bind within the “binding” protomer, whereas no ligands are seen in the “access” and “extrusion” protomers. In the “binding” protomer of AcrB-CU232, CU232 is also found within the distal drug-binding site. Amino acids within 4 Å of bound CU232 include F136, V139, Q151, F178, I277, A279, Y327, F610, V612, F615 and F628 (**Figure 3D**). Most of these 11 residues are hydrophobic and aromatic in nature, underscoring that this EPM analog recognition is largely governed by hydrophobic and aromatic interactions.

### Structure of AcrB-CU032

We also produced the AcrB-CU032 complex by incubating AcrB-nanodisc with CU032. The reconstituted sample led to a high-quality cryo-EM map, permitting us to solve the cryo-EM structure of the AcrB-CU032 complex to a resolution of 2.62 Å (**Table 2**, **Figure S4**). As with CU244 and CU232, the bound CU032 molecule is only found within the “binding” protomer, leaving the drug-binding sites of both the “access” and “extrusion” protomers unoccupied. The binding mode of CU032 within the “binding” protomer is similar to those of CU244 and CU232, where CU032 is anchored in the distal drug-binding pocket. However, it appears that AcrB utilizes a slightly different subset of residues to secure this modulator. Within 4 Å of bound CU032, residues F136, V139, F178, I277, A279, Y327, F610, V612, F615 and F628 participates in anchoring this ligand (**Figure 3E**). All 10 residues are hydrophobic and aromatic in nature, denoting that modulator binding is mostly controlled by hydrophobic and aromatic interactions.

### Structure of AcrB-EPM35

We also assembled the AcrB-EPM35 complex and solved its cryo-EM structure. The single-particle images led to a cryo-EM map at nominal resolution of 2.71 Å (**Table 2**, **Figure S5**). Like AcrB-CU244, AcrB-CU232 and AcrB-CU032, the trimeric AcrB-EPM35 complex is asymmetric in conformation and displays the “access”, “binding” and “extrusion” protomer structures. Pairwise superimpositions of the AcrB-CU244, AcrB-CU232, AcrB-CU032 and AcrB-EPM35 trimer provide r.m.s.d. values between 0.4 and 0.6 Å, suggesting that the structures of these four trimers are nearly identical to each other. The bound EPM35 molecule is only found within the “binding” protomer of AcrB, where its binding mode is very similar to those of CU244, CU232 and CU032. EPM35 largely resides at the distal drug-binding pocket of the periplasmic domain of the “binding” protomer. Within 4 Å of bound EPM35, residues F136, V139, F178, I277, A279, Y327, F610, V612, F615 and F628 are responsible for anchoring this modulator (**Figure 3F**). These 10 residues are identical to those involved in CU032 binding. Again, it is observed that AcrB contacts EPM35 via hydrophobic and aromatic interactions. No hydrogen bonds or significant electrostatic interactions are found to facilitate the binding.

### Proton Motive Force is not Dissipated by the EPM Analogs

The structural data suggest that the EPM analogs could be substrates of efflux pumps. However, the EPM analogs could also indirectly interfere with pump activity: their amphipathic nature suggests they could interact with membranes and damage the inner membrane barrier, which efflux pumps require for energy. We therefore monitored membrane integrity in the presence of EPM analogs by evaluating membrane voltage with mNeongreen-FtsZ (28–30). The FtsZ protein localizes to the bacterial septum based on membrane potential, and rapid delocalization of fluorescent FtsZ reporters occurs when voltage across the membrane is disrupted (31, 32). We monitored mNeongreen-FtsZ localization in a Gram-positive bacterium, *Bacillus subtills,* to eliminate the potentially confounding issue of the passage of EPM analogs through the outer membrane at different rates (24, 33–36). As expected, treatment of *B. subtilis* cells with the protonophore CCCP [100 μM] delocalized FtsZ (**Figure 4**) (31). We determined that CU032 and CU232, two of the compounds examined with AcrB by cryo-EM, had no significant effect on the size of the mNeongreen-FtsZ ring after 15 minutes of treatment at 150 μM. Likewise, CU187, the most active EPM analog in SAFIRE, and the EPM35 parent compound did not alter FtsZ localization. These data suggest that the EPM analogs do not indirectly inhibit efflux pumps by damaging membranes, dissipating proton motive force, and thereby generally depleting energy.

**Figure 4.**
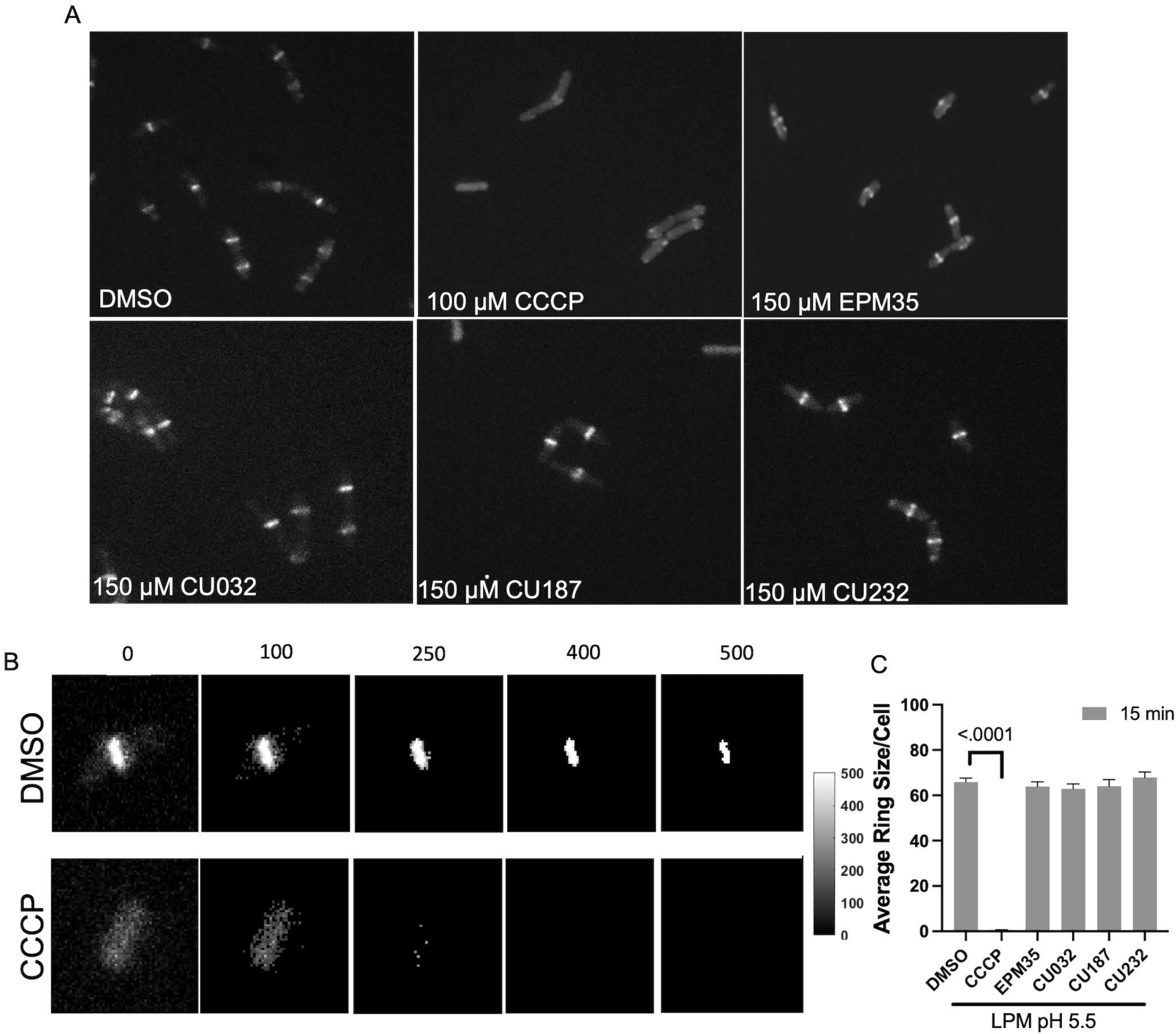
The EPM analogs minimally disrupt membrane voltage. A) **Micrographs of** *Bacillus subtilis* expressing mNeongreen-FtsZ were grown to mid log-phase in LPM 5.5. Samples were pipetted onto LPM 5.5 agar pads containing either DMSO, or with 150 μM of EPM35 or the EPM analog indicated for 15 minutes and imaged by confocal fluorescent microscopy. **B)** How FtsZ rings were quantified: overall intensity levels per pixel were quantified and a threshold of 400 was set, above which signal was measured to establish the FtsZ ring size within the cell. Examples of thresholds are shown. **C)** Cells exposed to EPM35 or EPM analogs were imaged at 15 after treatment, as indicated. Y axis is the average ring size in pixels per cell. For CCCP, the ring size was on average 1 pixel per cell. At least 50 cells from 3 biological replicates were compared for each sample. Error bars are SEM. *P* value of < 0.05 compared to DMSO calculated with a one-way ANOVA and a Tukey-Kramer posttest.

### AcrAB-TolC Likely Exports a Toxic Bacterial Metabolite Under Broth Conditions that Mimic the Phagosome Microenvironment

Given that the EPM analogs appear to directly inhibit AcrAB-TolC activity, we examined whether they interact with antibiotics to inhibit growth in media. Checkerboard assays are typically performed in Mueller Hinton Broth (MHB) (37, 38), but in this medium CU032 did not inhibit the growth of wild-type bacteria (**Figure S6**), prohibiting a full analysis of EPM-antibiotic interactions. However, CU032 prevented bacterial growth in LPM 5.5, a medium that mimics the microenvironment of the macrophage phagosome and can result in a leaky outer membrane (34) (**Figure 5A)**. LPM 5.5 is chemically defined, contains low phosphate, low magnesium, and is buffered to pH 5.5 (34, 39–41). Changing the pH of this medium to 7.0 abolished the CU032 inhibitory activity, which was bacteriostatic, not bactericidal (**Figure 5B).** It was further noted that mutant strains lacking *acrAB* or *tolC* grow poorly in LPM 5.5 compared to the corresponding wild-type (**Figure 5C-F)**. These observations suggest that in LPM 5.5, *S.* Typhimurium utilizes AcrAB-TolC to export a growth-inhibitory metabolite. The addition of CU032 or EPM35 to the medium of the *ΔacrAB* or *ΔtolC* mutant strains does not strongly inhibit growth compared to DMSO, suggesting that neither compound has a significant target beyond efflux pumps. In sum, the data support that LPM 5.5 is an appropriate medium for quantifying EPM-antibiotic interactions and show that in this medium, *S.* Typhimurium appears to accumulate an unknown, bacteriostatic metabolite(s) that is normally expelled by AcrAB-TolC.

**Figure 5.**
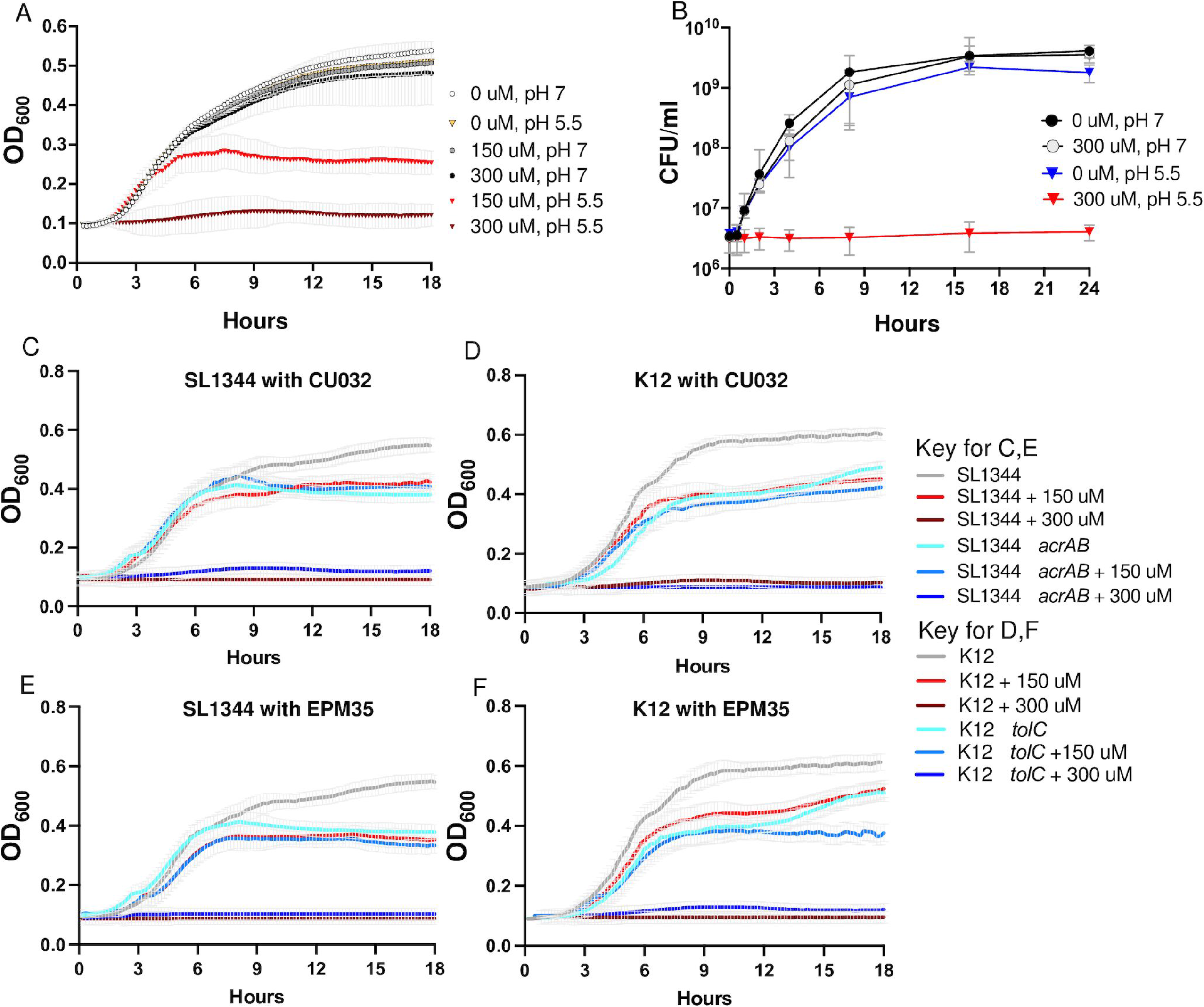
In LPM pH 5.5, *acrAB* and *tolC* are required for growth, and EPM analogs inhibit growth. **A, B)** OD_600_ and CFU/mL of *S.* Typhimurium in LPM 5.5 (at pH 5.5) or LPM 7 (at pH 7.0) in the presence of CU032. **C - F)** OD_600_ in LPM 5.5 of **C, E)** *S.* Typhimurium *ΔacrAB* mutant and wild-type strains, and **D, F)** *E. coli ΔtolC* mutant and wild-type strains with **C, D**) CU032 or **E, F**) EPM35. Mean and SEM of 3 biological replicates are shown for A-F. The wild-type and mutant strain controls are the same for C and E and for D and F.

### EPM Analogs Enhance the Potency of Ciprofloxacin and Erythromycin

To examine whether the EPM analogs interact additively or synergistically with antibiotics, checkerboard growth assays were carried out in LPM 5.5. We compared three antibiotics (ciprofloxacin, doxycycline, erythromycin) and three EPM analogs (CU032, CU187, CU232) and the EPM35 parent compound (**Figure 6A)**. Standard fractional inhibitory concentration index (FICI) analyses supported synergy between the EPM analogs and ciprofloxacin (**Figure 6B)**. Since FICI values cannot distinguish between synergy due to changes in potency (activity) versus efficacy (the extent of the effect) (42, 43), we used the MuSyC synergy algorithm to measure EPM-antibiotic synergy (44, 45). The data were well fit by the MuSyC equation, with an R2 of greater than 0.96 for all checkerboards (**Table S2**). Combining each of the EPM analogs with any one of the three antibiotics did not increase the maximal observed antibiotic efficacy by more than 7% (β_obs ≤ 0.07, **Figure 6C, Table S2**). Instead, all three EPMs increased the potency of each antibiotic by up to 72-fold (log_10_(α2) ≤ 1.87 and the increase was largest for ciprofloxacin, followed by erythromycin, then doxycycline (**Figure 6D, Table S2**). In contrast, the antibiotics did not increase the potency of the EPM analogs by more than 2-fold (log(α1) < 0.4). The EPM35 parent compound did not synergize with the antibiotics, indicating that the SAFIRE-based screening process identified compounds that are better at inhibiting efflux pumps than EPM35. Moreover, the consistency in the synergy of potency values across the antibiotics with different EPM analogs points towards a shared mechanism of action: the EPMs could modify the pharmacodynamics of the efflux process and prevent export of specific substrates through interactions with AcrB. We posit that the EPMs differentially reduce antibiotic export resulting in a higher effective intracellular concentration of the antibiotic.

**Figure 6.**
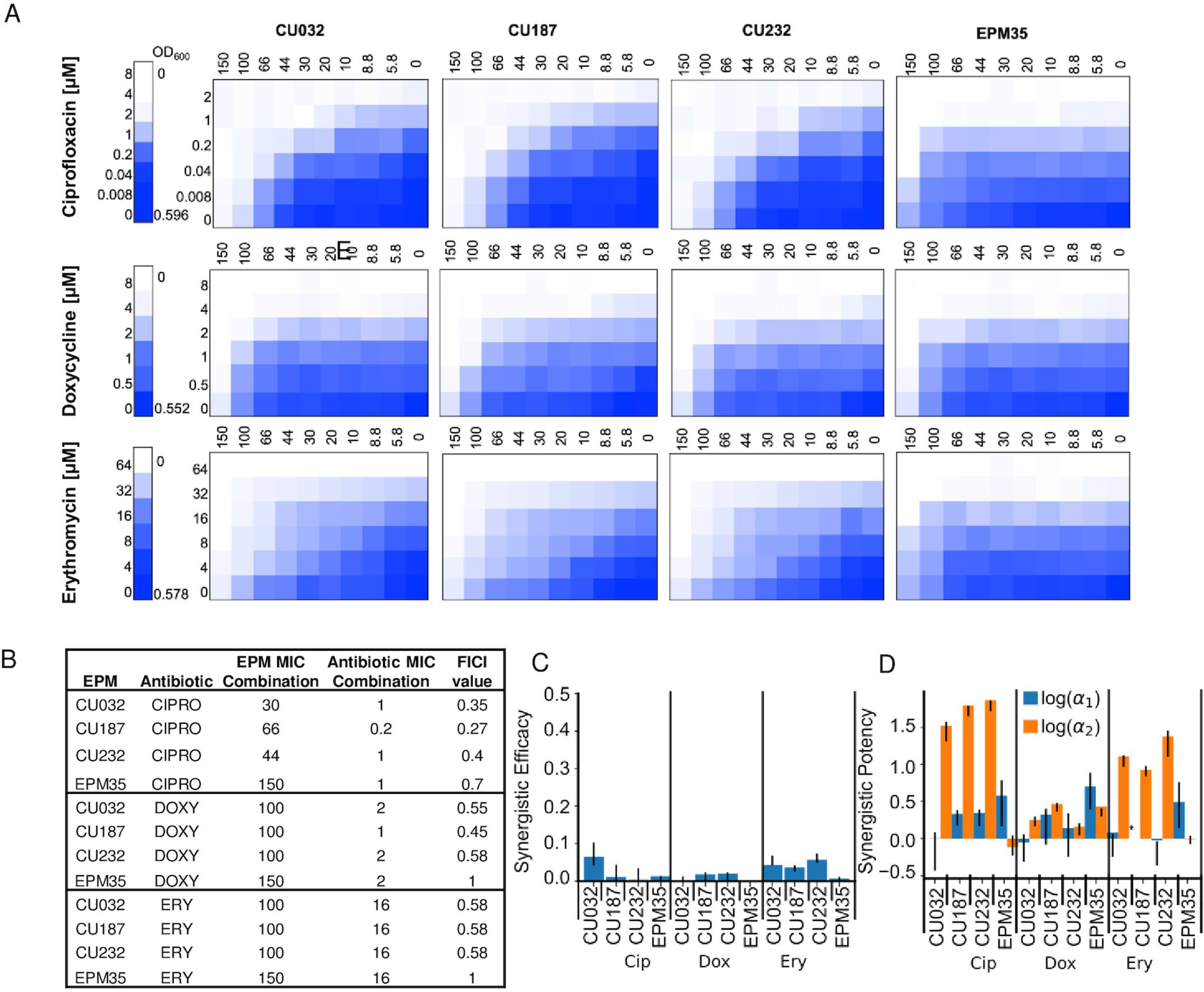
EPM analogs, but not the parent EPM35 compound, increase the potency of ciprofloxacin and erythromycin. **A)** Checkerboard assays with *S.* Typhimurium and three classes of clinical antibiotics. Date are means of at least three biological replicates. **B)** FICI calculations. **C, D)** MuSyC comparisons. **C)** β_abs_ values demonstrate little effect of EPM35 or the EPM analogs on antibiotic efficacy. **D)** Alpha1 (blue) represents the effect of the antibiotic on the compound, and alpha 2 (orange) the effect of the compound on the antibiotic.

## DISCUSSION

We set out to increase the potency and decrease the toxicity of a series previously described EPMs (18). Iterative medicinal chemistry and screening in the SAFIRE cell culture assay identified EPMs with 1,000-fold increases in activity, with toxicity below 50 μM. ITC and cryo-EM studies demonstrated that the EPM analogs specifically bind the AcrB inner membrane protein of the AcrAB-TolC efflux pump. The ITC-derived K_D_ of the EPM analogs was approximately 20-fold higher than that of the parent compound. However, all of these AcrB and EPM binding interactions are within the micromolar range, consistent with the binding affinities of AcrB with a variety of compounds (25). For instance, the *Klebsiella pneumoniae* AcrB multidrug efflux pump, which shares 91% protein sequence identity with AcrB from *E. coli*, distinctively contacts the antibiotic erythromycin with a K_D_ value of 14.4 ± 2.6 μM (46). This binding affinity is in good agreement with the strength of AcrB-EPM interactions revealed in this work. Within AcrB, the analogs occupy the hydrophobic substrate binding pocket, which is large and captures diverse molecules (13, 16, 25, 47). The EPMs interact with multiple amino acid residues in the pocket through hydrophobic bonds: no hydrogen bonding nor salt bridges were observed despite having the resolution required to detect them. These observations reveal that the EPMs may interfere with pump function in a manner that disrupts infection.

Despite their amphipathic structures, the EPMs do not appear to disrupt membranes. This is an important observation because dissipation of the proton motive force starves the cell for energy and indirectly reduces efflux. Delocalization of the septal protein FtsZ is a rapid and sensitive assay for PMF disruption (31, 32). We used a *B. subtilis* model system for this assay to avoid issues of compound exclusion from the cell membrane by an outer membrane barrier, an approach that could be useful for establishing whether other apparent EPMs damage membranes. While we worked with a MNeongreen labeled FtsZ protein for this purpose (48), other proteins also localize to the bacterial cell septum or membrane because they have a short alpha-helical region that associates with charged membranes (49, 50). Thus, a number of reporters (32) could be used to determine whether inhibitors of efflux adversely affect the proton gradient.

Efflux pumps are virulence determinants; they are not required for bacterial replication in standard laboratory media but are needed during infection in mammalian cells and/or animals (6, 51). In accordance with these prior observations, inhibitors of efflux pumps, including the EPM parent compounds, their analogs, and a series of promising pyranopyridines, do not prevent bacterial growth in standard microbiological media, specifically cation-adjusted MHB and/or lysogeny broth (LB) (18, 52, 53). We found that in LPM 5.5 medium, *acrAB* and *tolC* are required for *S.* Typhimurium and *E. coli* growth, respectively. LPM 5.5 was designed to mimic the acidified microenvironment of the macrophage phagosome (39–41) and potentiates the activity of antimicrobials (34, 54). Bacterial growth in LPM 5.5 could require *acrAB* and *tolC* because a bacteriostatic toxic metabolite(s) needs to be expelled by efflux pumps in this medium. This same potential metabolite(s) could accumulate in bacteria treated with the EPM analogs in LPM 5.5, explaining why the analogs inhibit growth under these conditions. For these reasons, LPM 5.5 medium could provide researchers with a future opportunity to identify an endogenous substrate(s) of AcrAB-TolC of relevance to the role of efflux pumps in virulence.

The medium LPM 5.5 mimics the macrophage phagosome in that it is acidic and has low levels of phosphate and magnesium. The latter are needed for outer membrane integrity: magnesium stabilizes LPS (55, 56), and phosphate plays a key structural role in LPS glycosidic linkages (57–60). Bacteria grown in LPM 5.5 have demonstrated increased outer membrane permeability compared to bacteria grown in standard media (34, 54). In LPM 5.5, the growth inhibitory activity of the EPM analogs could reflect that small molecules have improved access to AcrB and therefore prevent the export of a toxic metabolite at lower concentrations. Similarly, EPMs could potentiate antibiotics in LPM 5.5 due to improved antibiotic access to the bacterial cell. LB medium, like LPM 5.5, is deficient in divalent cations (61–63), suggesting that pyranopyridine efflux pump inhibitors act synergistically with antibiotics in LB (47) due to leakage of antibiotics and/or of the pump inhibitors across the outer membranes. Conditions within the macrophage phagosome also damage Gram-negative bacterial outer membranes, facilitate antibiotic access, and expose the pathogen to host-derived antimicrobials (64–66). Therefore, the EPMs could be particularly effective at reducing bacterial load in macrophages compared to in LPM 5.5 due to a combination of the accumulation of putative toxic bacterial metabolites and/or of host cell-autonomous defenses.

The use of the MuSyC synergy algorithm to analyze the checkerboard data from growth in LPM 5.5 enabled deeper probing of EPM-antibiotic interactions than standard FICI analyses (42, 43). MuSyC fits checkerboard data to a ten parameter, two-dimensional Hill equation. It requires a Hill-like dose–response curve for each single compound and distinguishes between synergy resulting from increases in potency, efficacy and/or cooperativity (44, 45). MuSyC revealed that the interaction of the EPM analogs and antibiotics occurs directionally: the EPMs increase the effective concentration of the antibiotics (synergy of potency), but the reverse does not occur. Furthermore, the EPMs do not increase the maximal extent to which antibiotics prevent growth (efficacy), instead they reduce the dose at which antibiotics inhibit growth (potency). The antibiotics in turn do not enhance the ability of the EPMs to prevent the export of a potential toxic metabolite(s). These observations support a causal relationship between the biochemical (i.e., AcrB-binding) and phenotypic activity (i.e., growth inhibition) of the EPMs. In other words, EPM binding to AcrB appears to directly interfere with the activity of the efflux pump.

Limitations of these studies include that LPM 5.5 does not fully mimic the macrophage phagosome microenvironment, which is dynamic and includes antimicrobial peptides, lysozyme, complement, and additional host antimicrobials and innate immune defenses recruited from vesicles and the cytosol (67–69). In addition, the EPM parent compounds or analogs could have unknown off-target activities. Data indicating that the parent compounds have bacterial targets beyond AcrB includes the observation that they reduce *ΔacrAB* mutant *S.* Typhimurium accumulation in macrophages below that of DMSO-treated controls (18). Possible alternative targets for the EPMs in macrophages include the multiple *acrB* orthologs encoded by Enterobacteriaceae (7, 14). It is also conceivable that in macrophages, the parent compounds or EPM analogs have a mammalian target(s). We conclude that the 1,000-fold increased activity of the EPM analogs in the SAFIRE macrophage assay relative to the parent EPMs could be due to a combination of factors, including increased access to bacteria caused by a weakened outer membrane barrier, the accumulation within intracellular bacteria of toxic efflux pump substrates, lysosomal trapping (35, 70), and possibly undefined off-target effects.

In summary, iterations of medicinal chemistry and a series of unconventional assays, including SAFIRE, cryo-EM, FtsZ localization, and checkerboards in LPM 5.5 medium, has improved upon and further characterized a family of EPMs. Specifically, these approaches enabled the generation of a set of related compounds that bind AcrB and appear to target efflux pumps in their capacity as virulence determinants, by preventing the export of an unknown toxic metabolite(s).

## METHODS

### Bacterial strains and media

*S*. *enterica* subsp. *enterica*, serovar Typhimurium strain SL1344 (71); *S*. Typhimurium strain *ΔacrAB* (ALR1257) in the SL1344 background (18); *S*. Typhimurium strain 14028s (ATCC); *S*. Typhimurium strain S10801, NR-22067 is a multidrug resistant isolate from a calf with sepsis (72) obtained through BEI resources, NIAID, NIH. *E*. *coli* (K-12 derivative BW25113 (wildtype)) (73); *E*. *coli* K-12 *ΔtolC* (JP313 *ΔtolC* (74), also called AD3644 and JLD1285). *Bacillus subtilis* Strain DK5092 (mNeonGreen-FtsZ) was constructed by DBK.

Bacteria were grown in Lysogeny Broth (LB) at 37°C (75, 76) unless otherwise stated. Cation-adjusted MHB was purchased from Sigma-Aldrich (90922). LPM 5.5 medium (5 mM KCl, 7.5 mM (NH_4_)2SO_4_, 0.5 mM K_2_SO_4_, 10 mM Glucose, 49 μM MgCl_2_, 337 μM PO_4_-, 0.05% CAS-Amino Acids, 80 mM MES. Adjust pH to 5.5 with 5M NaOH. Sterile Filter). Where stated, bacteria were grown with antibiotics: 30 μg/ml streptomycin.

### SAFIRE Screening for potency and toxicity

The initial screening of compounds in SAFIRE was performed at two concentrations, 5 and 25 μM, followed by dose response curves with a minimum of 8 concentrations per compound to determine the IC_50_. As previously described (18), RAW 264.7 macrophages (between passages 1 and 6) were grown in complete Dulbecco’s modified Eagle’s medium (DMEM) to approximately 70 to 90% confluence, scraped, washed, resuspended, diluted to 5 × 10^4^ in 100 μL, and seeded in black, 96-well, glass-bottomed plates (Brooks Life Sciences catalog number MGB096-1-2-LG-L) at 37°C with 5% CO_2_. Twenty-four hours later, bacteria grown overnight in LB were diluted into 50 μL PBS and added to a final concentration of 1 × 10^7^ *S.* Typhimurium SL1344 (*sifB*::*gfp*)/mL (77) for an approximate multiplicity of infection of 30 bacteria per macrophage cell. Forty-five minutes after bacterial addition, gentamicin (Sigma catalog number G1264) was added to a final concentration of 40 μg/mL to inhibit the growth of remaining extracellular bacteria. Two hours after infection, gentamicin-containing medium was removed and replaced with 200 μL fresh DMEM with compound or DMSO (Sigma catalog number 276855) to the stated final concentrations. At 17.5 h after infection, PBS containing MitoTracker Red CMXRos (Life Technologies catalog number M7512), a vital dye for mitochondrial electric potential, was added to a final concentration of 100 nM. At 18 h after infection, 16% paraformaldehyde was added to a final concentration of 4% and incubated at room temperature for 15 min. Cells were washed twice with PBS and stained for 20 min with 1 μM 4′,6-diamidino-2-phenylindole (DAPI), a vital dye for double stranded DNA, and stored in 90% glycerol in PBS until imaging. Samples were imaged in six fields of view per well using a semiautomated Yokogawa CellVoyager CV1000 confocal scanner system with a 20×, 0.75-numerical aperture (NA) objective. A MATLAB algorithm calculated bacterial accumulation (GFP fluorescence) within macrophages, as defined by DAPI and MitoTracker Red. GFP^+^ macrophage area was defined as the number of GFP-positive pixels per macrophage divided by the total number of pixels per macrophage, averaged across all macrophages in the field (34).

### Isothermal Titration Calorimetry (ITC)

AcrB protein was purified as described (25). Briefly, the AcrB protein contains a 4·His tag at the C-terminus and was overproduced in *E. coli* BL21-Gold (DE3) cells (Stratagene) using the plasmid derived from pSPORT1 (Invitrogen) (78). Cells were grown in 6 L of LB medium with 100 μg/ml ampicillin and disrupted with a French pressure cell. The membrane fraction was collected and washed twice with buffer containing 20 mM sodium phosphate (pH 7.2), 2 M KCl, 10% glycerol, 1 mM EDTA and 1 mM phenylmethanesulfonyl fluoride (PMSF), and once with 20 mM HEPES–NaOH buffer (pH 7.5) containing 1 mM PMSF. The membrane proteins were then solubilized in 1% (w/v) n-dodecylb-D-maltoside (DDM). Insoluble material was removed by ultracentrifugation at 370000 x g. The extracted protein was purified with Cu^2+^-affinity and G-200 sizing columns (13, 79). The purified AcrB protein was then concentrated to a final monomeric concentration of 10 μM in buffer containing 20 mM Na-HEPES (pH 7.5) and 0.05% DDM. Similar protein purification procedures were extensively used to elucidate structure-function of RND transporters, including AcrB (13, 16, 25), CusA (80, 81), MtrD (82, 83), CmeB (84), AdeB (85), HpnN (86) and MmpL3 (87). We also used these protein purification protocols for *in vitro* substrate transport study via the CusA transporter (81) and *in vitro* functional dynamics measurement of the CmeB transporter (84), indicating that these purified membrane proteins are fully functional *in vitro*.

Briefly, measurements were performed on a Microcal iTC200 (Malvern Panalytical) at 25°C. Before titration, the protein was dialyzed against buffer containing 20 mM Na-HEPES (pH 7.5), 0.05% n-dodecyl-µ-maltoside (DDM) and 5% DMSO (18). The Bradford assay was used to quantify protein concentration, which was adjusted to a final monomeric concentration of 10 µM. Ligand solution consisting of 100 µM EPM in the aforementioned buffer was prepared as the titrant. Both the protein and ligand samples were degassed before loading the samples. Two µL injections of the ligand were used for data collection. Injections occurred at intervals of 60 seconds and lasted for 4 seconds. Heat transfer (µcal/s) were measured as a function of elapsed time (s). The mean enthalpies measured from injection of the ligand in the buffer were subtracted from raw titration data before data analysis with ORIGIN software (MicroCal). Titration curves fitted with a non-linear regression fitting to the binding isotherm provided the equilibrium binding constant (*K_A_* = 1/*K_D_*) and enthalpy of binding (*ΔH*). Based on the values of *K_A_*, the change in free energy (*ΔG*) and entropy (*ΔS*) were calculated with the equation *ΔG* = -*RT* ln*K_A_* = *ΔH* - *TΔS*, where *T* is 2/3 K and *R* is 1.9872 cal/K per mol. Calorimetry trials were also carried out in the absence of AcrB using the same experimental conditions. No change in heat was observed in the injections throughout the experiment.

### Cryo-Electron Microscopy (Cryo-EM)

AcrB was purified from *E. coli* BL21(DE3)*ΔacrB*/pSPORTΩ*acrB* cells and reconstituted into lipidic nanodiscs. To assemble AcrB into nanodiscs, a mixture containing 10 μM AcrB, 30 μM membrane scaffold protein (MSP; 1E3D1), and 900 μM E. coli total extract lipid was incubated for 15 min at room temperature. After, 0.8 mg/ml prewashed Bio-Beads (Bio-Rad) was added. The resultant mixture was incubated for 1 h on ice, followed by overnight incubation at 4°C. The protein-nanodisc solution was filtered through 0.22-μm nitrocellulose filter tubes to remove the Bio-Beads. To separate free nanodiscs from AcrB-loaded nanodiscs, the filtered protein-nanodisc solution was purified using a Superose 6 column (GE Healthcare) equilibrated with 20 mM Tris-HCl (pH 7.5) and 100 mM NaCl. Fractions corresponding to the size of the trimeric AcrB-nanodisc complex were collected for cryo-EM (46). We then incubated 0.6 mg/mL AcrB-nanodisc with 10 μM EPM for 2 h and solved structure of the AcrB-EPM complex using single-particle cryo-electron microscopy (cryo-EM) to a resolution of 2.20 Å.

### Membrane voltage disruption assay (FtsZ delocalization)

To determine whether exposure to compound dislodges FtsZ from the cell septum, an overnight culture of *Bacillus subtilis* grown in LPM at pH 5.5 (LPM 5.5) expressing mNeonGreen-FtsZ (strain DK5092) was exposed to DMSO, CCCP [100 μM] or an EPM analog [150 μM] for 15 or 30 minutes, as indicated. Bacteria were immobilized on 1% agarose pads made with LPM under a coverslip. To set the threshold value for the FtsZ septal ring, mNeonGreen signal (excitation 488 nm, emission 525 nm) from at least 20 DMSO-treated cells was quantified using a CV1000 confocal microscope with a 100x oil, NA = 1.4 objective. A background-subtracted image was made by subtracting the median intensity from full original images. A signal threshold of ≥ 400 arbitrary units per pixel was set (Figure 4B). Then, we manually confirmed that mNeonGreen signal above threshold was at the center of the bacterial rod. We examined 50 cells per sample from at least 3 biological replicates (≥150 cells per condition). A MATLAB program (https://github.com/Betterton-Lab/FtsZ-Quantification) counted the number of pixels above threshold per cell and averaged this value across all cells for each condition and this value is reported in the graph (Figure 4C).

### Checkerboard Assays and Analyses

Overnight cation-adjusted MHB-grown cultures of SL1344 or SA10801 were diluted in MHB or LPM 5.5 to an optical density at 600 nm (OD_600_) of 0.01 and distributed into polystyrene 96-well flat-bottom plates. EPM analogs were added at concentrations up to 300 μM, near the limit of solubility, antibiotics ciprofloxacin, doxycycline or erythromycin was added up to a concentration of 2 μM, 2 μM and 8 μM respectively. The final DMSO concentration was at or below 2% in all wells. Plates were grown at 37°C with shaking for 18 hours and OD_600_ was monitored (BioTek Synergy H1, BioTek Eon). Each experiment was performed with biological triplicate or higher as indicated. Percent growth was determined by normalizing to OD_600_ reads in wells containing no D66 or antibiotic. Checkerboards in LPM 5.5 medium were performed with culture grown overnight in LB and diluted to an OD_600_ of 0.01 in LPM 5.5 and compounds/antibiotics were added as previously described. MuSyC analysis was pursued as previously described (44, 45). Briefly, the 2D Hill equation was fit to the normalized growth inhibition data from the checkerboards using a bounded non-linear least square regression algorithm. A Monte Carlo sampling method was used to compute the 95% confidence interval in the parameter fits from three biological replicates. The bounds on the parameters minimal and maximal efficacy of the antibiotics and EPMs (E_0_,E_max_) were [0.99,1.01] and [0,1], respectively (44, 45).

## ACKNOWLEDGEMENTS

We thank DB Kearns for providing the mNeongreen-FtsZ strain and for expert advice, C Morl for expert advice and editing, J. Orth for microscopy expertise, the Crestone Inc. team for medicinal chemistry, and G.L. Christensen and CA Ewing for experimental assistance, editing, and insightful comments.

## SUPPLEMENTAL MATERIAL

**Fig S1.**
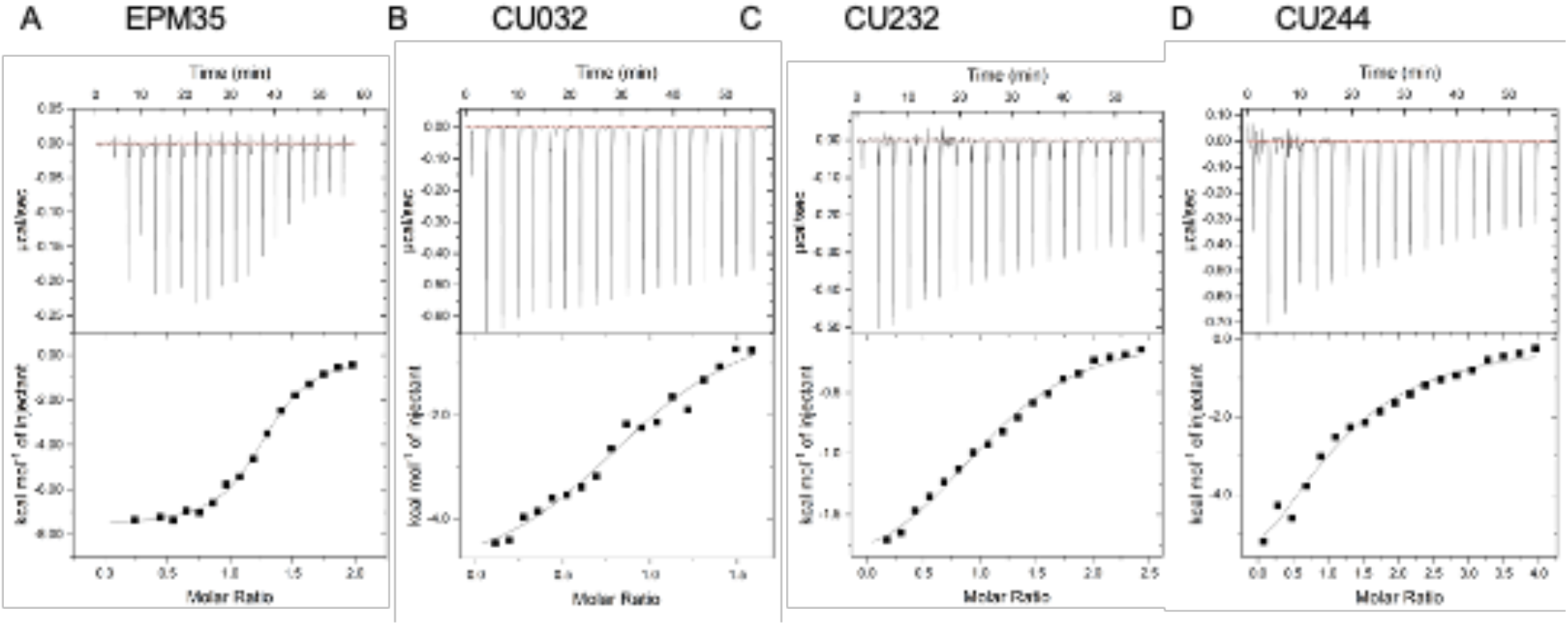
Binding of the EPM analogs to AcrB as shown by ITC. **A - D)** Representative ITC for the binding of EPM35 and three analogs to AcrB. Each peak in the upper panel corresponds to the injection of 2 μL of 100 μM of EPM in buffer containing 20 mM Na-HEPES (pH7.5), 0.05% DDM and 5% DMSO into the reaction containing 10 μM of monomeric AcrB in the same buffer. The lower panel shows the cumulative heat of reaction displayed as a function of injection number. The solid line is the least-square fit to the experimental data.

**Figure S2.**
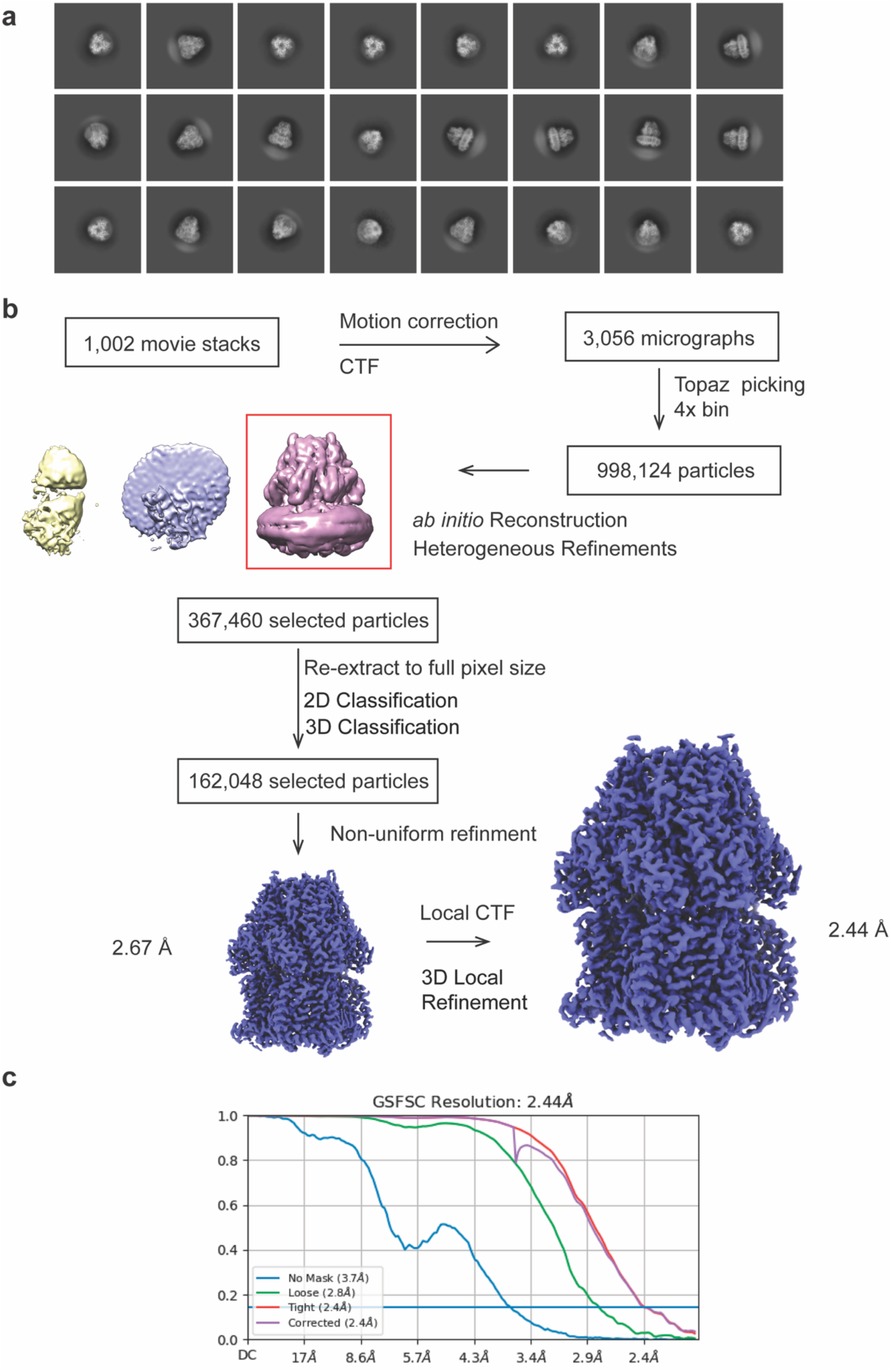
AcrB-CU244 data processing. **A)** Representative 2D classes of AcrB-CU244. **B)** Data processing workflow of AcrB-CU244. The side view density map of AcrB-CU244 is colored blue. (c) Gold-Standard Fourier shell correlation (GS-FSC) curves of AcrB-CU244, showing final resolution of 2.44 A.

**Figure S3.**
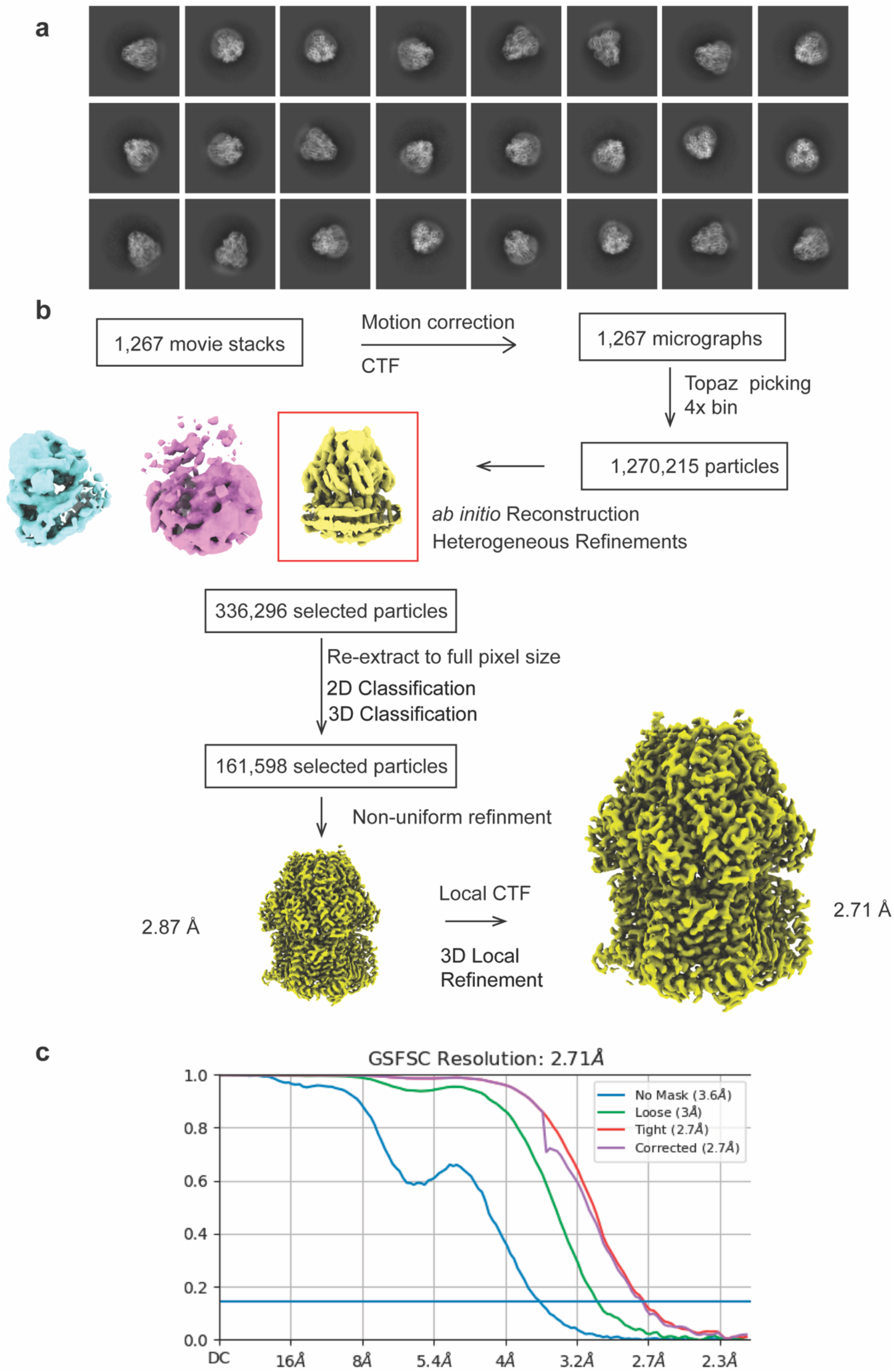
AcrB-CU232 data processing. **A)** Representative 2D classes of AcrB-CU232. **B)** Data processing workflow of AcrB-CU232. The side view density map of AcrB-CU232 is colored yellow. (c) Gold-Standard Fourier shell correlation (GS-FSC) curves of AcrB-CU232, showing final resolution of 2.71 A.

**Figure S4.**
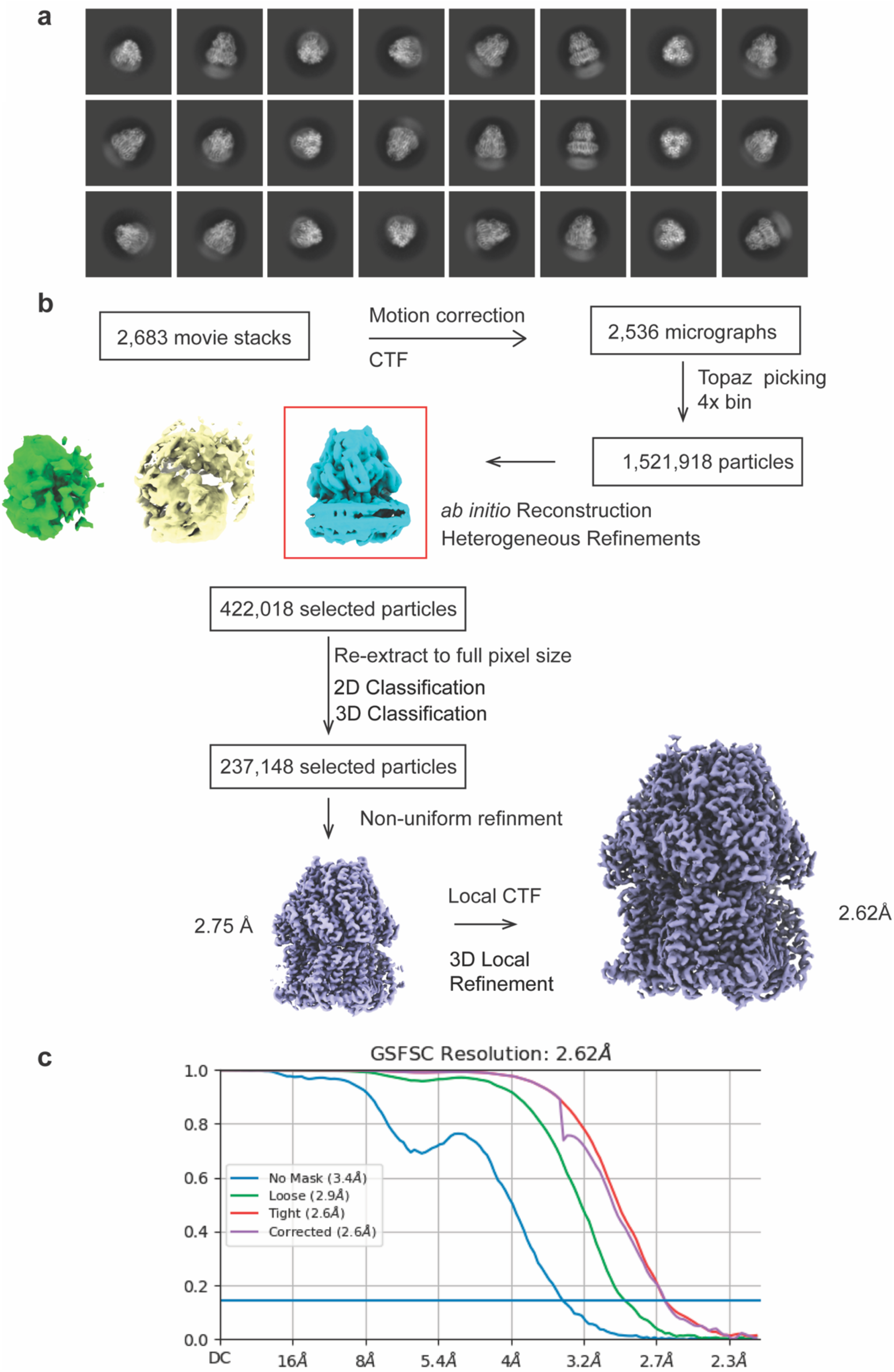
AcrB-CU032 data processing. **A)** Representative 2D classes of AcrB-CU032. **B)** Data processing workflow of AcrB-CU032. The side view density map of AcrB-CU032 is colored slate. (c) Gold-Standard Fourier shell correlation (GS-FSC) curves of AcrB-CU032, showing final resolution of 2.62 A.

**Figure S5.**
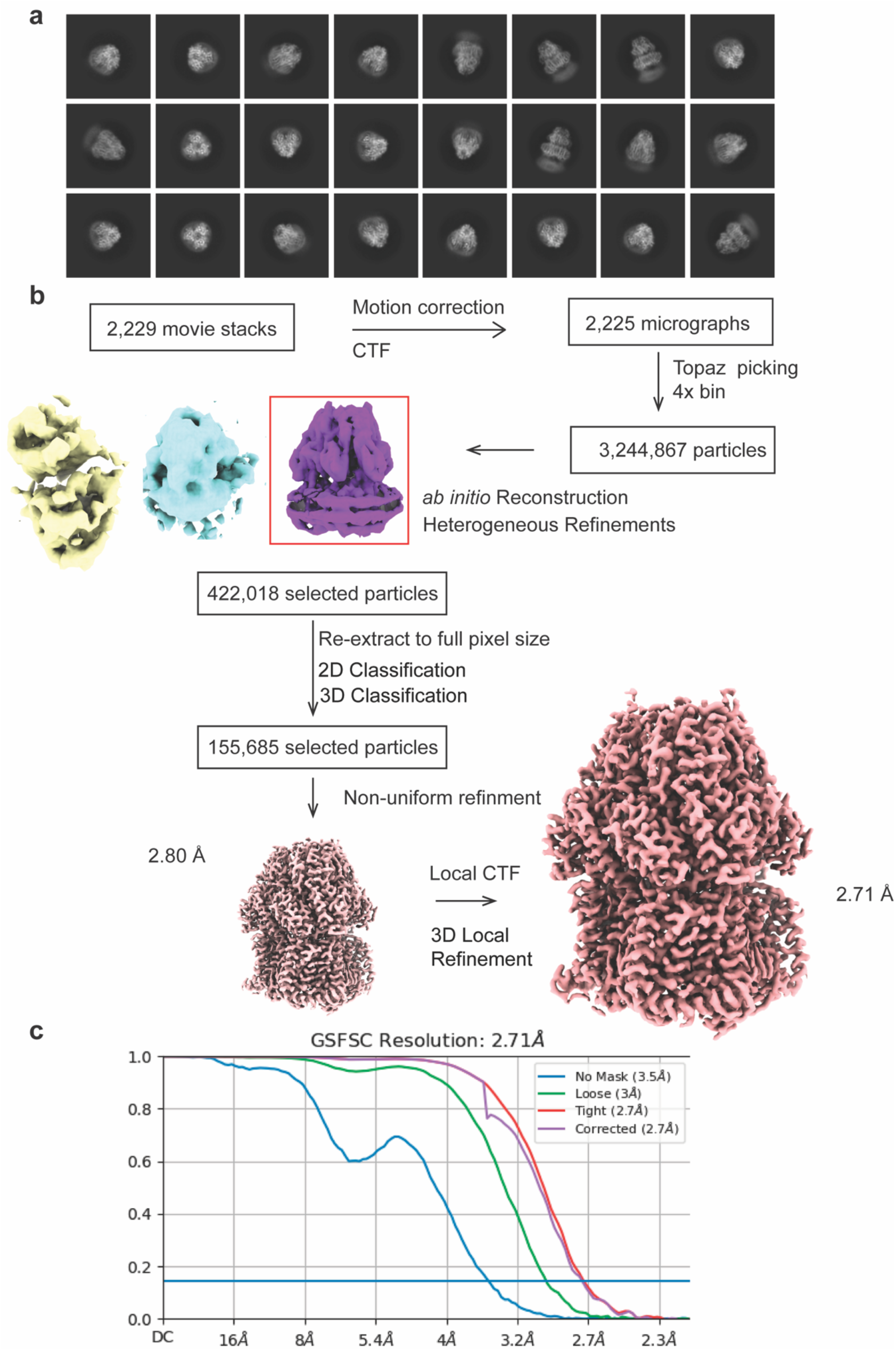
AcrB-EPM35 data processing. **A)** Representative 2D classes of AcrB-EPM35. **B)** Data processing workflow of AcrB-CU035. The side view density map of AcrB-EPM35 is colored pink. (c) Gold-Standard Fourier shell correlation (GS-FSC) curves of AcrB-EPM35, showing final resolution of 2.71 A.

**Figure S6.**
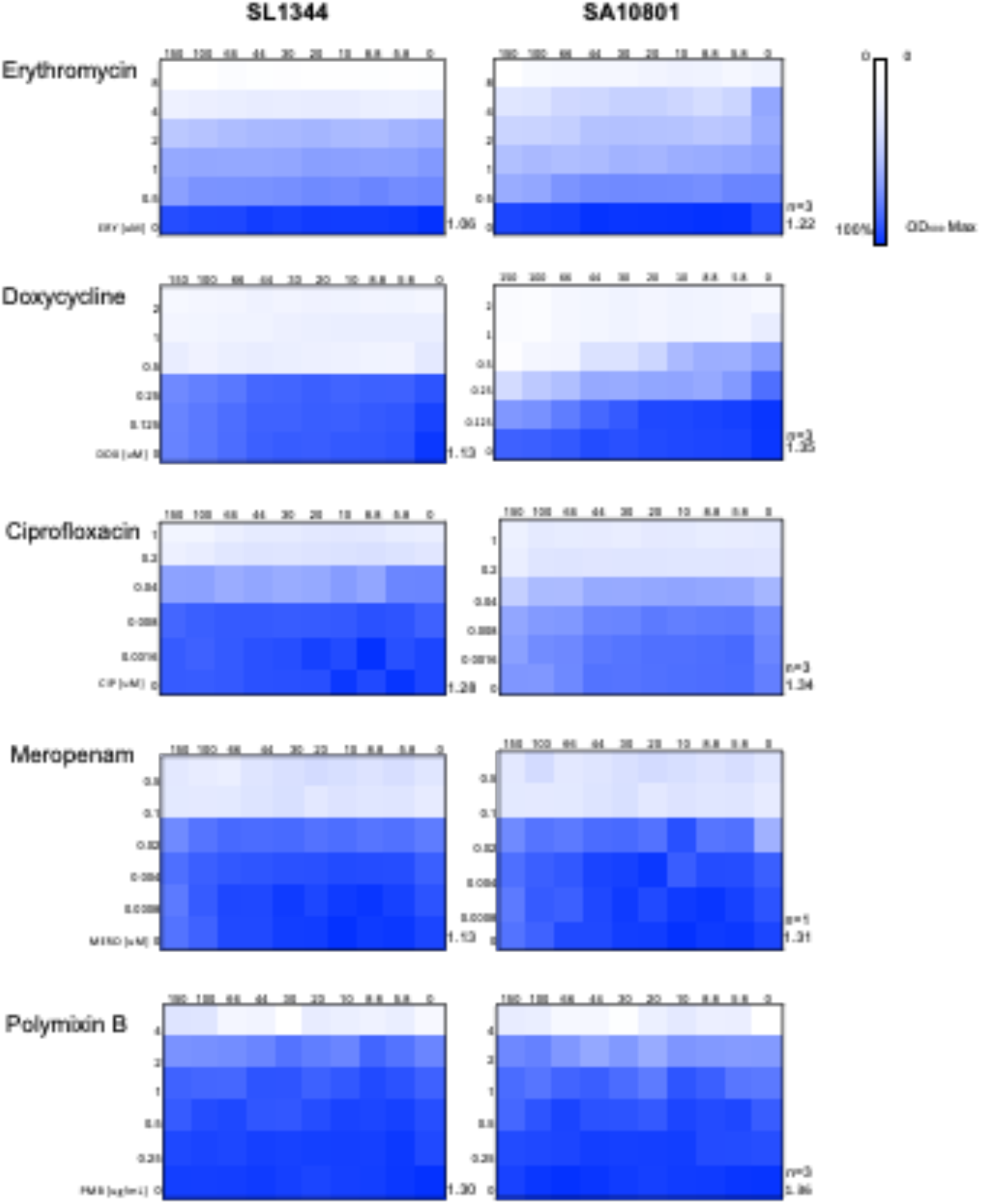
CU032 did not inhibit the growth of *S. enterica* in cation-adjusted MHB. Checkerboard Assays with CU032 and four classes of clinical antibiotics or the antimicrobial peptide polymyxin B. Two virulent *S. enterica* clinical isolates (S1344 and SA10801) are shown.

---

**Table S1** Analog Testing in SAFIRE (see downloadable xls file)

**Table S2.** MuSyC synergy fit values.

| Antibiotic | EPM | Fit $R^2$ | $B_{obs}$ [95% CI]<sup>1</sup> | $\log_{10}(\alpha_1)$ [95% CI]<sup>2</sup> | $\log_{10}(\alpha_2)$ [95% CI]<sup>3</sup> |
| --- | --- | --- | --- | --- | --- |
| Ciprofloxacin | CU032 | 0.96 | 0.07 [0.05,0.1 ] | 0.0 [-0.42,0.07] | 1.52 [1.32,1.56] |
|  | CU187 | 0.98 | 0.01 [-0.01,0.04] | 0.33 [0.19,0.37] | 1.79 [1.66,1.78] |
|  | CU232 | 0.98 | 0.0 [-0.02,0.03] | 0.34 [0.18,0.38] | 1.87 [1.73,1.85] |
|  | EPM35 | 0.98 | 0.01 [-0.01,0.01] | 0.58 [0.17,0.77] | -0.12 [-0.22,0.03] |
| Doxycycline | CU032 | 0.98 | 0.0 [-0.01,0.01] | -0.06 [-0.3,0.04] | 0.25 [0.17,0.28] |
|  | CU187 | 0.97 | 0.02 [-0.0,0.02] | 0.32 [-0.07,0.39] | 0.46 [0.37,0.47] |
|  | CU232 | 0.97 | 0.02 [-0.01,0.02] | 0.14 [-0.23,0.33] | 0.16 [0.06,0.19] |
|  | EPM35 | 0.98 | -0.01 [-0.04,-0.01] | 0.7 [0.41,0.87] | 0.43 [0.31,0.39] |
| Erythromycin | CU032 | 0.98 | 0.04 [0.04,0.07] | 0.08 [-0.23,0.05] | 1.11 [0.98,1.11] |
|  | CU187 | 0.99 | 0.04 [0.03,0.04] | nan [-4.0,-0.82] | 0.93 [0.85,0.96] |
|  | CU232 | 0.98 | 0.06 [0.05,0.07] | -0.02 [-0.35,-0.05] | 1.38 [1.11,1.44] |
|  | EPM35 | 0.99 | 0.01 [-0.01,0.01] | 0.49 [0.15,0.74] | -0.01 [-0.06,0.03] |

<sup>1</sup> $B_{obs}$ is the proportion increase in maximal efficacy of the antibiotics at maximum tested dose of EPMs.
<sup>2</sup> $\log_{10}(\alpha_1)$ is the log-fold-change in potency of EPMs due to the antibiotics. Shown in brackets are the 95% confidence interval (CI) for the MuSyC fits.
<sup>3</sup> $\log_{10}(\alpha_2)$ is the log-fold-change in potency of the antibiotics due to the EPMs.

## REFERENCES

1. Witwinowski J, Sartori-Rupp A, Taib N, Pende N, Tham TN, Poppleton D, Ghigo J-M, Beloin C, Gribaldo S. 2022. An ancient divide in outer membrane tethering systems in bacteria suggests a mechanism for the diderm-to-monoderm transition. Nat Microbiol 7:411–422.

2. Zgurskaya HI, Rybenkov VV. 2020. Permeability barriers of Gram-negative pathogens. Ann N Y Acad Sci 1459:5–18.

3. Taib N, Megrian D, Witwinowski J, Adam P, Poppleton D, Borrel G, Beloin C, Gribaldo S. 2020. Genome-wide analysis of the Firmicutes illuminates the diderm/monoderm transition. Nat Ecol Evol 4:1661–1672.

4. Bertani B, Ruiz N. 2018. Function and Biogenesis of Lipopolysaccharides. EcoSal Plus 8.

5. Silhavy TJ, Kahne D, Walker S. 2010. The bacterial cell envelope. Cold Spring Harb Perspect Biol 2:a000414.

6. Li X-Z, Plésiat P, Nikaido H. 2015. The challenge of efflux-mediated antibiotic resistance in Gram-negative bacteria. Clin Microbiol Rev 28:337–418.

7. Nishino K, Latifi T, Groisman EA. 2006. Virulence and drug resistance roles of multidrug efflux systems of Salmonella enterica serovar Typhimurium. Mol Microbiol 59:126–141.

8. Bogomolnaya LM, Andrews KD, Talamantes M, Maple A, Ragoza Y, Vazquez-Torres A, Andrews-Polymenis H. 2013. The ABC-type efflux pump MacAB protects Salmonella enterica serovar typhimurium from oxidative stress. mBio 4:e00630–00613.

9. Buckley AM, Webber MA, Cooles S, Randall LP, La Ragione RM, Woodward MJ, Piddock LJV. 2006. The AcrAB-TolC efflux system of Salmonella enterica serovar Typhimurium plays a role in pathogenesis. Cell Microbiol 8:847–856.

10. Lacroix FJ, Cloeckaert A, Grépinet O, Pinault C, Popoff MY, Waxin H, Pardon P. 1996. Salmonella typhimurium acrB-like gene: identification and role in resistance to biliary salts and detergents and in murine infection. FEMS Microbiol Lett 135:161–167.

11. Theuretzbacher U. 2017. Global antimicrobial resistance in Gram-negative pathogens and clinical need. Curr Opin Microbiol 39:106–112.

12. Aron Z, Opperman TJ. 2018. The hydrophobic trap-the Achilles heel of RND efflux pumps. Res Microbiol 169:393–400.

13. Yu EW, McDermott G, Zgurskaya HI, Nikaido H, Koshland DE. 2003. Structural basis of multiple drug-binding capacity of the AcrB multidrug efflux pump. Science 300:976–980.

14. Nishino K, Nikaido E, Yamaguchi A. 2009. Regulation and physiological function of multidrug efflux pumps in Escherichia coli and Salmonella. Biochim Biophys Acta 1794:834–843.

15. Zgurskaya HI, Malloci G, Chandar B, Vargiu AV, Ruggerone P. 2021. Bacterial efflux transporters’ polyspecificity - a gift and a curse? Curr Opin Microbiol 61:115–123.

16. Yu EW, Aires JR, McDermott G, Nikaido H. 2005. A periplasmic drug-binding site of the AcrB multidrug efflux pump: a crystallographic and site-directed mutagenesis study. J Bacteriol 187:6804–6815.

17. Klenotic PA, Morgan CE, Yu EW. 2021. Cryo-EM as a tool to study bacterial efflux systems and the membrane proteome. Fac Rev 10:24.

18. Reens AL, Crooks AL, Su C-C, Nagy TA, Reens DL, Podoll JD, Edwards ME, Yu EW, Detweiler CS. 2018. A cell-based infection assay identifies efflux pump modulators that reduce bacterial intracellular load. PLoS Pathog 14:e1007115.

19. Detweiler CS. 2020. Infection-based chemical screens uncover host-pathogen interactions. Curr Opin Microbiol 54:43–50.

20. Nakashima R, Sakurai K, Yamasaki S, Nishino K, Yamaguchi A. 2011. Structures of the multidrug exporter AcrB reveal a proximal multisite drug-binding pocket. Nature 480:565– 569.

21. Kinana AD, Vargiu AV, Nikaido H. 2013. Some ligands enhance the efflux of other ligands by the Escherichia coli multidrug pump AcrB. Biochemistry 52:8342–8351.

22. Rieg S, Huth A, Kalbacher H, Kern WV. 2009. Resistance against antimicrobial peptides is independent of Escherichia coli AcrAB, Pseudomonas aeruginosa MexAB and Staphylococcus aureus NorA efflux pumps. Int J Antimicrob Agents 33:174–176.

23. Liu J, Huang Z, Ruan B, Wang H, Chen M, Rehman S, Wu P. 2020. Quantitative proteomic analysis reveals the mechanisms of polymyxin B toxicity to Escherichia coli. Chemosphere 259:127449.

24. Dombach JL, Quintana JLJ, Nagy TA, Wan C, Crooks AL, Yu H, Su C-C, Yu EW, Shen J, Detweiler CS. 2020. A small molecule that mitigates bacterial infection disrupts Gram-negative cell membranes and is inhibited by cholesterol and neutral lipids. PLoS Pathog 16:e1009119.

25. Su C-C, Nikaido H, Yu EW. 2007. Ligand-transporter interaction in the AcrB multidrug efflux pump determined by fluorescence polarization assay. FEBS Lett 581:4972–4976.

26. Murakami S, Nakashima R, Yamashita E, Matsumoto T, Yamaguchi A. 2006. Crystal structures of a multidrug transporter reveal a functionally rotating mechanism. Nature 443:173–179.

27. Vargiu AV, Nikaido H. 2012. Multidrug binding properties of the AcrB efflux pump characterized by molecular dynamics simulations. Proc Natl Acad Sci 109:20637–20642.

28. Bisson-Filho AW, Hsu Y-P, Squyres GR, Kuru E, Wu F, Jukes C, Sun Y, Dekker C, Holden S, VanNieuwenhze MS, Brun YV, Garner EC. 2017. Treadmilling by FtsZ filaments drives peptidoglycan synthesis and bacterial cell division. Science 355:739–743.

29. Yu Y, Zhou J, Gueiros-Filho FJ, Kearns DB, Jacobson SC. 2021. Noc Corrals Migration of FtsZ Protofilaments during Cytokinesis in Bacillus subtilis. mBio 12:e02964–20.

30. Oshiro RT, Rajendren S, Hundley HA, Kearns DB. 2019. Robust Stoichiometry of FliW-CsrA Governs Flagellin Homeostasis and Cytoplasmic Organization in Bacillus subtilis. mBio 10:e00533–19.

31. Strahl H, Hamoen LW. 2010. Membrane potential is important for bacterial cell division. Proc Natl Acad Sci U S A 107:12281–12286.

32. Schäfer A-B, Wenzel M. 2020. A How-To Guide for Mode of Action Analysis of Antimicrobial Peptides. Front Cell Infect Microbiol 10:540898.

33. Dombach JL, Quintana JL, Allgood SC, Nagy TA, Gustafson DL, Detweiler CS. 2022. A small molecule that disrupts S. Typhimurium membrane voltage without cell lysis reduces bacterial colonization of mice. PLoS Pathog 18:e1010606.

34. Nagy TA, Crooks AL, Quintana JLJ, Detweiler CS. 2020. Clofazimine Reduces the Survival of Salmonella enterica in Macrophages and Mice. ACS Infect Dis 6:1238–1249.

35. Villanueva JA, Crooks AL, Nagy TA, Quintana JLJ, Dalebroux ZD, Detweiler CS. 2022. Salmonella enterica Infections Are Disrupted by Two Small Molecules That Accumulate within Phagosomes and Differentially Damage Bacterial Inner Membranes. mBio e0179022.

36. Ude J, Tripathi V, Buyck JM, Söderholm S, Cunrath O, Fanous J, Claudi B, Egli A, Schleberger C, Hiller S, Bumann D. 2021. Outer membrane permeability: Antimicrobials and diverse nutrients bypass porins in Pseudomonas aeruginosa. Proc Natl Acad Sci U S A 118:e2107644118.

37. Mueller JH, Hinton J. 1941. A Protein-Free Medium for Primary Isolation of the Gonococcus and Meningococcus. Proc Soc Exp Biol Med 48:330–333.

38. Nizet V. 2017. The Accidental Orthodoxy of Drs. Mueller and Hinton. EBioMedicine 22:26– 27.

39. Coombes BK, Brown NF, Valdez Y, Brumell JH, Finlay BB. 2004. Expression and secretion of Salmonella pathogenicity island-2 virulence genes in response to acidification exhibit differential requirements of a functional type III secretion apparatus and SsaL. J Biol Chem 279:49804–49815.

40. Beuzón CR, Banks G, Deiwick J, Hensel M, Holden DW. 1999. pH-dependent secretion of SseB, a product of the SPI-2 type III secretion system of Salmonella typhimurium. Mol Microbiol 33:806–816.

41. Deiwick J, Nikolaus T, Erdogan S, Hensel M. 1999. Environmental regulation of Salmonella pathogenicity island 2 gene expression. Mol Microbiol 31:1759–1773.

42. Odds FC. 2003. Synergy, antagonism, and what the chequerboard puts between them. J Antimicrob Chemother 52:1–1.

43. Meyer CT, Wooten DJ, Lopez CF, Quaranta V. 2020. Charting the Fragmented Landscape of Drug Synergy. Trends Pharmacol Sci 41:266–280.

44. Wooten DJ, Meyer CT, Lubbock ALR, Quaranta V, Lopez CF. 2021. MuSyC is a consensus framework that unifies multi-drug synergy metrics for combinatorial drug discovery. Nat Commun 12:4607.

45. Meyer CT, Wooten DJ, Paudel BB, Bauer J, Hardeman KN, Westover D, Lovly CM, Harris LA, Tyson DR, Quaranta V. 2019. Quantifying Drug Combination Synergy along Potency and Efficacy Axes. Cell Syst 8:97–108.e16.

46. Zhang Z, Morgan CE, Bonomo RA, Yu EW. 2023. Cryo-EM Structures of the Klebsiella pneumoniae AcrB Multidrug Efflux Pump. mBio e0065923.

47. Sjuts H, Vargiu AV, Kwasny SM, Nguyen ST, Kim H-S, Ding X, Ornik AR, Ruggerone P, Bowlin TL, Nikaido H, Pos KM, Opperman TJ. 2016. Molecular basis for inhibition of AcrB multidrug efflux pump by novel and powerful pyranopyridine derivatives. Proc Natl Acad Sci U S A 113:3509–3514.

48. Patrick JE, Kearns DB. 2008. MinJ (YvjD) is a topological determinant of cell division in Bacillus subtilis. Mol Microbiol 70:1166–1179.

49. Szeto TH, Rowland SL, Habrukowich CL, King GF. 2003. The MinD membrane targeting sequence is a transplantable lipid-binding helix. J Biol Chem 278:40050–40056.

50. Pichoff S, Lutkenhaus J. 2005. Tethering the Z ring to the membrane through a conserved membrane targeting sequence in FtsA. Mol Microbiol 55:1722–1734.

51. Baba T, Ara T, Hasegawa M, Takai Y, Okumura Y, Baba M, Datsenko KA, Tomita M, Wanner BL, Mori H. 2006. Construction of Escherichia coli K-12 in-frame, single-gene knockout mutants: the Keio collection. Mol Syst Biol 2:2006.0008.

52. Nguyen ST, Kwasny SM, Ding X, Cardinale SC, McCarthy CT, Kim H-S, Nikaido H, Peet NP, Williams JD, Bowlin TL, Opperman TJ. 2015. Structure-activity relationships of a novel pyranopyridine series of Gram-negative bacterial efflux pump inhibitors. Bioorg Med Chem 23:2024–2034.

53. Opperman TJ, Kwasny SM, Kim H-S, Nguyen ST, Houseweart C, D’Souza S, Walker GC, Peet NP, Nikaido H, Bowlin TL. 2014. Characterization of a novel pyranopyridine inhibitor of the AcrAB efflux pump of Escherichia coli. Antimicrob Agents Chemother 58:722–733.

54. Ellis MJ, Tsai CN, Johnson JW, French S, Elhenawy W, Porwollik S, Andrews-Polymenis H, McClelland M, Magolan J, Coombes BK, Brown ED. 2019. A macrophage-based screen identifies antibacterial compounds selective for intracellular Salmonella Typhimurium. Nat Commun 10:197.

55. Ellison RT, LaForce FM, Giehl TJ, Boose DS, Dunn BE. 1990. Lactoferrin and transferrin damage of the Gram-negative outer membrane is modulated by Ca2+ and Mg2+. Microbiology, 136:1437–1446.

56. Coughlin RT, Tonsager S, McGroarty EJ. 1983. Quantitation of metal cations bound to membranes and extracted lipopolysaccharide of Escherichia coli. Biochemistry 22:2002– 2007.

57. Rosner MR, Khorana HG, Satterthwait AC. 1979. The structure of lipopolysaccharide from a heptose-less mutant of Escherichia coli K-12. II. The application of 31P NMR spectroscopy. J Biol Chem 254:5818–5825.

58. Mühlradt PF, Wray V, Lehmann V. 1977. A 31P-nuclear-magnetic-resonance study of the phosphate groups in lipopolysaccharide and lipid A from Salmonella. Eur J Biochem 81:193–203.

59. Lehmann V, Rupprecht E. 1977. Microheterogeneity in lipid A demonstrated by a new intermediate in the biosynthesis of 3-deozy-D-manno-octulosonic-acid--lipid A. Eur J Biochem 81:443–452.

60. Rick PD, Fung LW, Ho C, Osborn MJ. 1977. Lipid A mutants of Salmonella typhimurium. Purification and characterization of a lipid A precursor produced by a mutant in 3-deoxy-D-mannooctulosonate-8-phosphate synthetase. J Biol Chem 252:4904–4912.

61. Nikaido H. The Limitations of LB Medium. Small Things Consid. https://schaechter.asmblog.org/schaechter/2009/11/the-limitations-of-lb-medium.html. Retrieved 15 January 2021.

62. Wee S, Wilkinson BJ. 1988. Increased outer membrane ornithine-containing lipid and lysozyme penetrability of Paracoccus denitrificans grown in a complex medium deficient in divalent cations. J Bacteriol 170:3283–3286.

63. Papp-Wallace KM, Maguire ME. 2008. Magnesium Transport and Magnesium Homeostasis. EcoSal Plus 3.

64. Heesterbeek DAC, Muts RM, van Hensbergen VP, de Saint Aulaire P, Wennekes T, Bardoel BW, van Sorge NM, Rooijakkers SHM. 2021. Outer membrane permeabilization by the membrane attack complex sensitizes Gram-negative bacteria to antimicrobial proteins in serum and phagocytes. PLoS Pathog 17:e1009227.

65. Coers J. 2017. Sweet host revenge: Galectins and GBPs join forces at broken membranes. Cell Microbiol 19.

66. Berti A, Rose W, Nizet V, Sakoulas G. 2020. Antibiotics and Innate Immunity: A Cooperative Effort Toward the Successful Treatment of Infections. Open Forum Infect Dis 7:ofaa302.

67. Kutsch M, Coers J. 2021. Human guanylate binding proteins: nanomachines orchestrating host defense. FEBS J 288:5826–5849.

68. Doorduijn DJ, Rooijakkers SHM, Heesterbeek DAC. 2019. How the Membrane Attack Complex Damages the Bacterial Cell Envelope and Kills Gram-Negative Bacteria. BioEssays News Rev Mol Cell Dev Biol 41:e1900074.

69. Pucciarelli MG, García-Del Portillo F. 2017. Salmonella Intracellular Lifestyles and Their Impact on Host-to-Host Transmission. Microbiol Spectr 5.

70. Goldman SDB, Funk RS, Rajewski RA, Krise JP. 2009. Mechanisms of amine accumulation in, and egress from, lysosomes. Bioanalysis 1:1445–1459.

71. Hoiseth SK, Stocker B a. D. 1981. Aromatic-dependent Salmonella typhimurium are non-virulent and effective as live vaccines. Nature 291:238–239.

72. Daniels JB, Call DR, Hancock D, Sischo WM, Baker K, Besser TE. 2009. Role of Ceftiofur in Selection and Dissemination of blaCMY-2-Mediated Cephalosporin Resistance in Salmonella enterica and Commensal Escherichia coli Isolates from Cattle. Appl Environ Microbiol 75:3648–3655.

73. Grenier F, Matteau D, Baby V, Rodrigue S. 2014. Complete Genome Sequence of Escherichia coli BW25113. Genome Announc 2:e01038–14.

74. Montaño ET, Nideffer JF, Sugie J, Enustun E, Shapiro AB, Tsunemoto H, Derman AI, Pogliano K, Pogliano J. 2021. Bacterial Cytological Profiling Identifies Rhodanine-Containing PAINS Analogs as Specific Inhibitors of Escherichia coli Thymidylate Kinase In Vivo. J Bacteriol 203:e0010521.

75. Bertani G. 1951. Studies on lysogenesis. I. The mode of phage liberation by lysogenic Escherichia coli. J Bacteriol 62:293–300.

76. Bertani G. 2004. Lysogeny at Mid-Twentieth Century: P1, P2, and Other Experimental Systems. J Bacteriol 186:595–600.

77. Rollenhagen C, Sörensen M, Rizos K, Hurvitz R, Bumann D. 2004. Antigen selection based on expression levels during infection facilitates vaccine development for an intracellular pathogen. Proc Natl Acad Sci U S A 101:8739–8744.

78. Takatsuka Y, Nikaido H. 2006. Threonine-978 in the transmembrane segment of the multidrug efflux pump AcrB of Escherichia coli is crucial for drug transport as a probable component of the proton relay network. J Bacteriol 188:7284–7289.

79. Zgurskaya HI, Nikaido H. 1999. Bypassing the periplasm: Reconstitution of the AcrAB multidrug efflux pump of Escherichia coli. Proc Natl Acad Sci 96:7190–7195.

80. Long F, Su C-C, Zimmermann MT, Boyken SE, Rajashankar KR, Jernigan RL, Yu EW. 2010. Crystal structures of the CusA efflux pump suggest methionine-mediated metal transport. Nature 467:484–488.

81. Su C-C, Long F, Zimmermann MT, Rajashankar KR, Jernigan RL, Yu EW. 2011. Crystal structure of the CusBA heavy-metal efflux complex of Escherichia coli. Nature 470:558– 562.

82. Bolla JR, Su C-C, Do SV, Radhakrishnan A, Kumar N, Long F, Chou T-H, Delmar JA, Lei H-T, Rajashankar KR, Shafer WM, Yu EW. 2014. Crystal structure of the Neisseria gonorrhoeae MtrD inner membrane multidrug efflux pump. PloS One 9:e97903.

83. Lyu M, Moseng MA, Reimche JL, Holley CL, Dhulipala V, Su C-C, Shafer WM, Yu EW. 2020. Cryo-EM Structures of a Gonococcal Multidrug Efflux Pump Illuminate a Mechanism of Drug Recognition and Resistance. mBio 11.

84. Su C-C, Yin L, Kumar N, Dai L, Radhakrishnan A, Bolla JR, Lei H-T, Chou T-H, Delmar JA, Rajashankar KR, Zhang Q, Shin Y-K, Yu EW. 2017. Structures and transport dynamics of a Campylobacter jejuni multidrug efflux pump. Nat Commun 8:171.

85. Su C-C, Morgan CE, Kambakam S, Rajavel M, Scott H, Huang W, Emerson CC, Taylor DJ, Stewart PL, Bonomo RA, Yu EW. 2019. Cryo-Electron Microscopy Structure of an Acinetobacter baumannii Multidrug Efflux Pump. mBio 10.

86. Kumar N, Su C-C, Chou T-H, Radhakrishnan A, Delmar JA, Rajashankar KR, Yu EW. 2017. Crystal structures of the Burkholderia multivorans hopanoid transporter HpnN. Proc Natl Acad Sci U S A 114:6557–6562.

87. Su C-C, Klenotic PA, Bolla JR, Purdy GE, Robinson CV, Yu EW. 2019. MmpL3 is a lipid transporter that binds trehalose monomycolate and phosphatidylethanolamine. Proc Natl Acad Sci U S A 116:11241–11246.

